# Spatial specificity of the functional gradient echo and spin echo BOLD signal across cortical depth at 7 T

**DOI:** 10.1101/2025.07.28.667324

**Authors:** Daniel Haenelt, Robert Trampel, Denis Chaimow, Amir Shmuel, Martin I. Sereno, Nikolaus Weiskopf

**Affiliations:** Department of Neurophysics, Max Planck Institute for Human Cognitive and Brain Sciences, 04103 Leipzig, Germany; International Max Planck Research School on Neuroscience of Communication: Function, Structure, and Plasticity, 04103 Leipzig, Germany; McConnell Brain Imaging Centre, Montreal Neurological Institute, Departments of Neurology, Neurosurgery, Physiology and Biomedical Engineering, McGill University, Montreal, QC H3A 2B4, Canada; Department of Psychology, College of Sciences, San Diego State University, San Diego, CA 92182; Felix Bloch Institute for Solid State Physics, Faculty of Physics and Earth Sciences, Leipzig University, 04103 Leipzig, Germany; Wellcome Centre for Human Neuroimaging, Institute of Neurology, University College London, London WC1N 3AR, UK

## Abstract

Functional magnetic resonance imaging (fMRI) at high magnetic field strengths (≥7 T) is a promising technique to study the functioning of the human brain at the spatial scale of cortical columns and layers. However, measurements most often rely on the blood oxygenation level dependent (BOLD) response sampled with a gradient echo (GE) sequence, which is known to be most sensitive to macrovascular contributions that limit their effective spatial resolution. Alternatively, a spin echo (SE) sequence can be used to increase the weighting toward the microvasculature and, therefore, the location of neuronal activation. In addition, due to the heterogeneous structure of the cortical cerebrovascular system, the effective spatial resolution can change across cortical depth. For high-resolution fMRI applications, it is hence important to know how much the effective spatial resolution varies across cortical depth. In this study, we used flickering rotating wedge stimuli to induce traveling waves with varying spatial frequencies in the retinotopically organized primary visual cortex (V1), which allowed us to infer the modulation transfer function (MTF) of the BOLD response that characterizes the spatial specificity of the measured signal. We acquired GE- and SE-BOLD data at 7 T and compared the MTF between acquisition techniques at different cortical depths. Although we observe a small but consistent increase in spatial specificity with SE-BOLD, both acquisition techniques exhibit a similar decrease in spatial specificity across cortical depth, with lower specificity toward the pial surface. This finding highlights the influence of macrovascular contributions on both imaging techniques, which must be carefully considered when interpreting results from high-resolution fMRI studies.

## Introduction

Functional magnetic resonance imaging (fMRI) is an indispensable non-invasive imaging technique for the study of human brain processes *in vivo* (Glover, 2011) and, therefore, a valuable tool for cognitive sciences and neuroscience in general (Poldrack, 2012). At high magnetic field strengths (7 T and above), high-resolution functional imaging with nominal resolutions reaching into the sub-millimeter regime is possible (Kim and Uğurbil, 2003; Uğurbil, 2021). Given that the cortical thickness of humans ranges on average from about 2 to 4 mm (Fischl and Dale, 2000), high-resolution fMRI allows the mapping of features at the mesoscopic scale like cortical columns (Cheng, Waggoner, and Tanaka, 2001; Yacoub et al., 2007; Nasr, Polimeni, and Tootell, 2016; Haenelt et al., 2023) and cortical layers, or laminae (Huber et al., 2017; Huber et al., 2021; Hollander et al., 2021; Iamshchinina et al., 2021). Cortical columns are defined as groups of neurons that share response properties across cortical depth (Mountcastle, 1957; Hubel and Wiesel, 1962) and are often viewed as the fundamental building block of the cortex (Mountcastle, 1997). The human cortex is further comprised of several layers that can be differentiated by their cell composition (Brodmann, 1909) and connectivity pattern (Felleman and Van Essen, 1991). The possibility to disentangle columnar and laminar features with fMRI, therefore, has the potential to study the mesoscopic regime of cortical organization, i.e., the local circuitry of the human brain (Douglas and Martin, 1991; Yang et al., 2021).

However, different mechanisms lead to a loss of spatial precision in fMRI and reduce the effective resolution that can be achieved, such as motion and processing steps during data analysis (Wang et al., 2022), geometric distortions (Jezzard, 2012), data sampling in the presence of 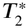 decay (Huber et al., 2015; Chaimow and Shmuel, 2016), use of the partial Fourier acquisition technique (Huber et al., 2015; Chaimow and Shmuel, 2016), and ultimately constraints by the underlying biology (Engel, Glover, and Wandell, 1997; Parkes et al., 2005; Chaimow et al., 2018). Since most fMRI acquisitions rely on the blood oxygenation level dependent (BOLD) contrast, which is an indirect measure of neuronal activity that exploits the hemodynamic response following neuronal activation (Buxton, 2013), biological constraints in terms of the underlying vascular architecture and dynamics (draining veins) represent the major limitation for the spatial specificity of the measured fMRI signal (Turner, 2002).

These limitations become particularly relevant in high-resolution fMRI, as the spatial specificity of the measured signal depends strongly on the underlying vascular architecture, which is characterized by a highly complex and spatially heterogeneous morphology (Duvernoy, Delon, and Vannson, 1981; Havlicek and Uludağ, 2020; Uludağ and Havlicek, 2021). From a network of arteries at the pial surface of the cerebral cortex, diving arteries branch off and supply the parenchyma with oxygenated blood. Deoxygenated blood is delivered back to the pial surface via ascending veins and drains into the pial veins (macrovascular compartments). Diving arteries and ascending veins are connected by a dense and randomly oriented capillary network (microvascular compartment). This organization generally leads to a microvascular system that is more homogeneously distributed across cortical depth compared to the macrovascular compartments (Weber et al., 2008).

The distinction between these vascular compartments is important for understanding the spatial specificity of the BOLD signal, as their contributions to the measured signal differ. BOLD signal contributions are generally divided into intra- and extravascular compartments, which can be described independently due to the slow exchange of water relative to the duration of fMRI image acquisition (Uludağ, Müller-Bierl, and Uğurbil, 2009). The gradient echo (GE) BOLD contrast that is most often used in fMRI, is known to be most sensitive to the intra- and extravascular signal contributions from macrovascular compartments (ascending veins, pial veins) (Boxerman et al., 1995; Uludağ, Müller-Bierl, and Uğurbil, 2009). Due to the organization of the macrovascular system, the GE-BOLD signal will show a typical signal increase across cortical depth toward the pial surface (Polimeni et al., 2010) and loss in spatial resolution (Fracasso, Dumoulin, and Petridou, 2021). This is mainly caused by the drainage of deoxygenated blood from deeper cortical depth (Markuerkiaga,

Barth, and Norris, 2016) and by dephasing around pial veins, which also manifests itself on the empirically found orientation dependence of 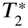-weighted signals on cortical orientation relative to the main magnetic field (Viessmann et al., 2019; Ramadan et al., 2025).

Alternatively, a spin echo (SE) BOLD contrast at high magnetic field strengths can be used (Yacoub et al., 2007). Using higher magnetic fields has the advantage of decreasing intravascular signal contributions due to the shortening of blood *T*_2_ (Thulborn et al., 1982; Yacoub et al., 2001). Furthermore, the application of a spin echo refocuses extravascular signal contributions from around larger draining veins (Boxerman et al., 1995), leading to a stronger weighting to the microvasculature, and therefore an increase of the effective spatial resolution. However, acquiring a pure spin-echo signal at high magnetic fields remains challenging because the long readout duration associated with fast data sampling methods such as echo-planar imaging (EPI) introduces 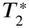-related contamination from macrovascular compartments into the SE-BOLD signal, limiting the achievable increase in spatial specificity.

In this study, we analyzed the effective resolution of GE- and SE-BOLD at different cortical depths by estimating the BOLD modulation transfer function (MTF). The MTF is a measure used to characterize an imaging system’s ability to faithfully resolve fine details of an imaged object and, therefore, the spatial resolution capabilities of the imaging system (Steckner, Drost, and Prato, 1994). More precisely, it describes the decrease of the signal amplitude as a function of spatial frequency of the imaged object. The Fourier transform of the MTF is the point spread function (PSF), which represents the response of an imaging system to a point source. The PSF’s width, typically defined by the full width at half maximum (FWHM), can be used to characterize the achievable effective spatial resolution.

Since the drainage of venous blood by ascending veins and pial veins is unidirectional, it is expected that the spatial specificity of the BOLD signal differs in radial (along the surface normal of the cortex) and tangential (perpendicular to surface normal) directions. Although our analysis is done across cortical depths, the estimated MTF in this study will only characterize the spatial spread in tangential direction at a specific cortical depth.

To estimate the MTF, we used a classic phase-encoded paradigm (Sereno et al., 1995; Engel, Glover, and Wandell, 1997) to induce traveling waves of cortical activation with different spatial wavelengths in retinotopically organized cortical areas (V1, V2, … ), such as the primary visual cortex (V1). We developed an MTF model that was incorporated in a hierarchical Bayesian regression framework based on Markov Chain Monte Carlo (MCMC) sampling to estimate the parameters of interest.

Similar experiments were already performed to estimate the spatial precision of GE-BOLD and SE-BOLD at lower field strengths (Engel, Glover, and Wandell, 1997; Parkes et al., 2005). A recent study also examined the PSF across cortical depth of GE-BOLD by exploiting the columnar organization of the secondary visual cortex (V2) (Fracasso, Dumoulin, and Petridou, 2021). In the current study, we show for the first time an estimation of the SE-BOLD PSF across cortical depth at 7 T, which allows a comprehensive comparison of spatial precision between GE- and SE-BOLD using echo-planar imaging (EPI) readouts.

## Methods

### Participants

Six healthy participants (2 females; age = 27.33 ± 4.15, mean ± standard deviation in years) gave written informed consent to participate in this study. The study was approved by the local ethics committee of the University of Leipzig. All participants had normal or corrected-to-normal visual acuity.

### General procedure

Each participant was invited five times for measurements in a 7 T scanner. The first session was used to acquire a high-resolution anatomical reference scan and retinotopy data. An additional functional time series without task (GE-BOLD) was measured to aid between-session registration and allow for estimating voxels with significant venous contributions. In the remaining four sessions (2× GE-BOLD, 2× SE-BOLD), we acquired functional data while using rotating wedges for stimulation (multi-polar sessions) from which we estimated the BOLD modulation transfer function (*MT F*_*BOLD*_). A subset of the retinotopy data was already used in another experiment (Movahedian Attar et al., 2020) but was independently processed for this study.

### Experimental design

Visual stimuli were presented using an LCD projector (Sanyo PLC-XT20L with custom-built focusing objective lens, refresh rate: 60 Hz, pixel resolution: 1024 × 768) that was positioned inside the magnet room. The projector was placed inside a custom-built Faraday cage to minimize interferences with the MR scanner. Stimuli were projected onto a rear-projection screen mounted above the participant’s chest inside the bore and were viewed through a mirror attached to the head coil. Black felt was put around the screen, and we turned off all lights during experiments to decrease the amount of scattered light reaching the participant’s eyes. The setup allowed the visual stimulation of around 22^°^× 13^°^(width × height) visual angle.

#### Multi-polar measurements

Participants were invited to four functional imaging sessions with GE- and SE-BOLD data acquired in two independent sessions each. In each session, seven functional runs were acquired, each lasting 528 s, resulting in a total functional acquisition time of approximately 62 min per session. For visual stimulation, we used a classical phase-encoded paradigm (Sereno et al., 1995; Engel, Glover, and Wandell, 1997) to induce traveling waves in retinotopically organized cortical areas. A flickering (4 Hz) black-and-white radial checkerboard restricted to a number of anticlockwise rotating wedges (number of wedges: 2, 4, …, 14) on a gray background with mean luminance 44 cd/m^2^ was shown in separate runs (see ***Figure 1***). Thus, the visual stimulus differed between runs within each session only in the number of wedges presented. The order of runs with different wedge numbers, and hence different spatial frequencies, was pseudo-randomized for each session. We chose a rotating wedge stimulus instead of inducing traveling waves in eccentricity direction (e.g., expanding ring stimulus) to circumvent the need to consider the significant changes in cortical representation due to cortical magnification (Horton and Hoyt, 1991). 10.5 stimulation cycles were presented per run, and the rotation speed was adjusted to maintain a constant stimulation period of 48 s despite the different number of wedges. Each run started and ended with 12 s of baseline, during which only a gray background was shown. During the runs, participants were instructed to fixate on a central point without performing an explicit task.

**Figure 1.**
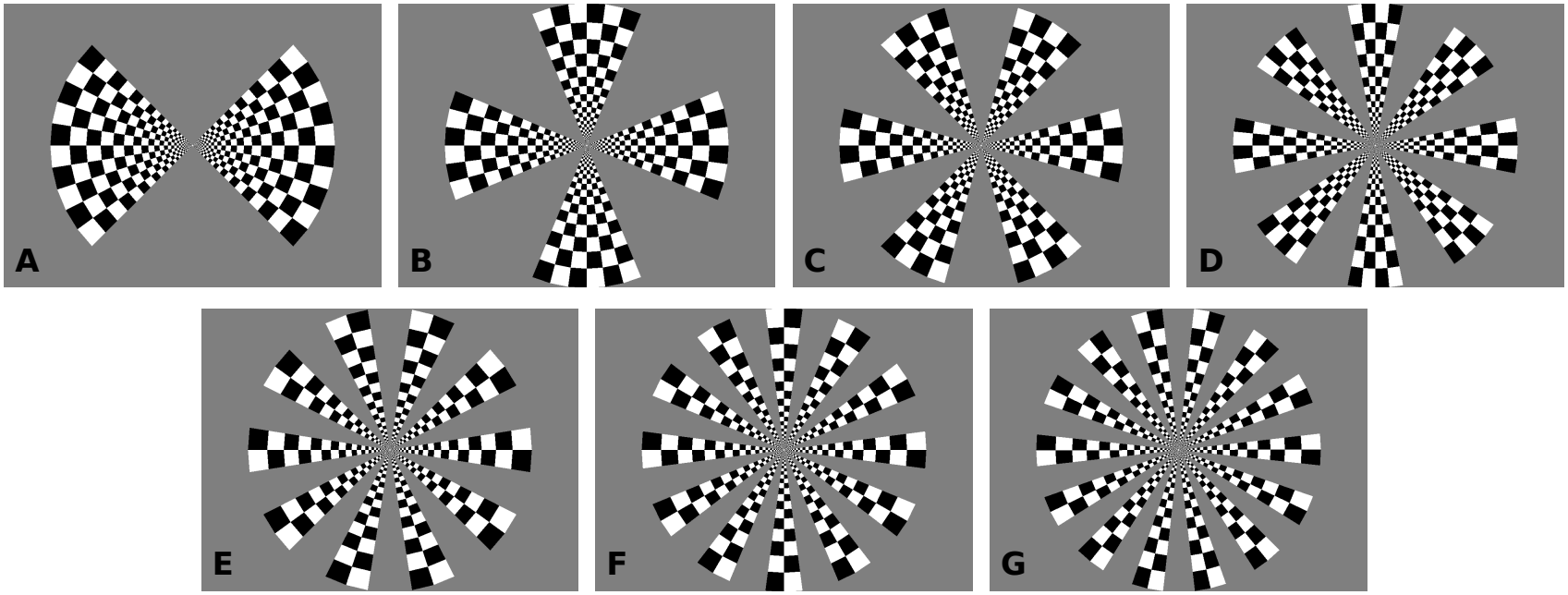
Illustration of visual stimuli. To create traveling waves of neuronal activity in retinotopically organized cortex at different spatial frequencies, we used flickering wedge stimuli with varying numbers of wedges that slowly rotated in counterclockwise direction. Different numbers of wedges {2, 4, …, 14} shown in **A–G** were used in separate runs.

#### Retinotopy measurements

We acquired retinotopy data from four runs to locate V1 and for the estimation of the length of iso-eccentricity lines that are needed to model the MTF in units of cycles per mm of cortex. For mapping of polar angle and eccentricity, we used the same flickering checkerboard restricted to a clockwise/anticlockwise wedge (angle: 30^°^, period: 64 s) and expanding/contracting ring (period: 32 s), respectively, presented in separate runs. 8.25 cycles were shown in each run.

### Imaging

All data were acquired with a 7 T whole-body MR scanner (MAGNETOM 7 T, Siemens Healthineers, Erlangen, Germany) equipped with SC72 body gradients (maximum gradient strength: 70 mT/m; maximum slew rate: 200 mT/m/s). For radio frequency signal transmission and reception, we used a single-channel transmit/32-channel receive head coil (Nova Medical, Wilmington, DE, USA). The transmit voltage was calibrated for optimal excitation over the occipital lobe by acquisition of a low-resolution transmit field map at the beginning of each scanning session using a sequence that eploits the ratio of consecutive spin and stimulated echoes (Siemens WIP-658).

For GE- and SE-BOLD functional measurements, we used a 2D single-shot EPI sequence (Feinberg et al., 2010; Moeller et al., 2010). We imaged an oblique-coronal slab positioned over the occipital lobe, which is illustrated for a representative participant in ***Figure 2***. Care was taken to cover the posterior part of the calcarine sulcus in all functional imaging sessions. Across sessions, we ensured that the slab position was consistently centered on the same anatomical location. For both acquisition techniques, we used the following parameters: nominal voxel size = 0.8 mm isotropic, repetition time (TR) = 3000 ms, field of view (FOV) = 148 × 148 mm^2^, readout bandwidth (rBW) = 1182 Hz/px, interleaved slice acquisition order. Partially parallel acquisition (GRAPPA) = 3 (Griswold et al., 2002) and partial Fourier = 6/8 with zero-filling of unsampled *k*-space lines were used for acceleration in the EPI phase-encoding direction, which resulted in an echo train length of around 46 ms with an effective echo spacing of 0.33 ms.

**Figure 2.**
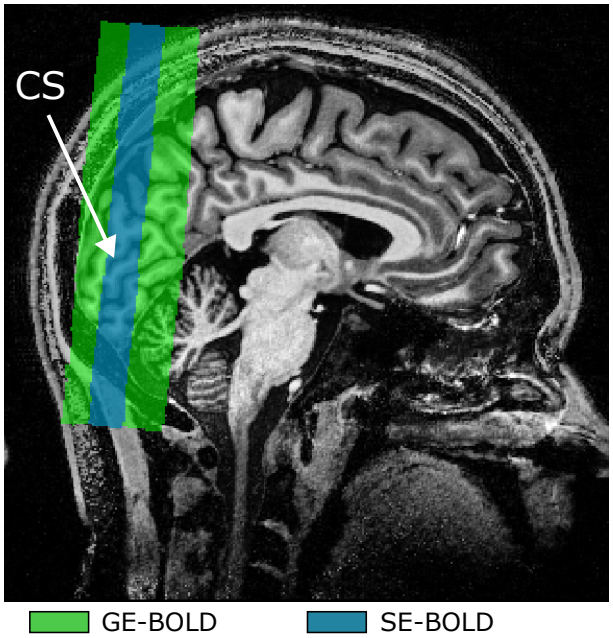
Illustration of slab positioning. The spatial coverage of GE-BOLD (green box) and SE-BOLD (blue box) acquisitions is overlaid on a *T*_1_-weighted anatomical image for one session of a representative participant (subject 6) in sagittal view. Importantly, the center slices of both acquisitions were explicitly matched and covered regions around the posterior calcarine sulcus (white arrow). Note the reduced spatial coverage of the SE-BOLD acquisition.

For GE-BOLD, the echo time (TE) was set to 24 ms, and the Ernst angle for the gray matter compartment was used as flip angle (77^°^). We could cover the whole stimulated area of V1 with 50 slices with GE-BOLD. For SE-BOLD, the number of slices varied due to specific absorption rate (SAR) constraints. We always used the maximum allowed number of slices that differed between subjects and sessions and was in the range of 18–32 slices. The echo time (TE) was set to 38 ms. For retinotopy measurements, we used a slightly modified GE-BOLD protocol with nominal voxel size = 1.0 mm isotropic, TR = 2000 ms, TE = 21 ms, FA = 68^°^, rBW = 1164 Hz/px, 40 slices.

For anatomical reference, we acquired a whole-brain anatomy using a 3D *T*_1_-weighted MP2RAGE sequence (Marques et al., 2010) with the following parameters: voxel size = 0.7 mm isotropic, TR = 5000 ms, TE = 2.45 ms, inversion times (TI1/TI2) = 900 ms/2750 ms with FA = 5^°^/3^°^, respectively, FOV = 224 × 224 × 168 mm^3^ (read × phase × partition), rBW = 250 Hz/px, partial Fourier = 6/8, and GRAPPA = 2 (primary phase-encoding direction; outer loop). From both inversion times, a uniform T1-weighted image (UNI) was created by the online image reconstruction on the scanner.

Protocols of all acquisitions are publicly available (https://osf.io/rzd3c/).

### Data preprocessing

Functional time series from multi-polar measurements were first corrected for different slice timings by voxel-wise temporal interpolation onto a common time grid using 3drefit from AFNI (19.1.05, https://afni.nimh.nih.gov/) (Cox, 1996). Within-run motion was corrected using SPM12 (v6906, https://www.fil.ion.ucl.ac.uk/spm/) with Matlab R2019b (MathWorks, Natick, MA, USA). Motion-corrected time series were then highpass filtered (cutoff frequency: 1/144 Hz) and converted to percent signal change by division by their temporal mean. We discarded the first half stimulus cycle, which left ten cycles for analysis. The amplitude at stimulation frequency *c*_*s*_ was computed by a voxel-wise Fourier analysis.

Functional time series from retinotopy measurements were similarly preprocessed. Cutoff frequencies for the highpass filter of polar angle and eccentricity runs were set to 1/192 Hz and 1/96 Hz, respectively. Using custom-built Matlab scripts, we applied standard population receptive field (pRF) modeling (Dumoulin and Wandell, 2008). Population receptive fields were modeled as isotropic 2D Gaussians with center position (*x, y*) and spread (*σ*) in units of degrees of visual angle. We estimated these parameters and the coefficient of determination (*r*^2^) for each voxel, which was then input to the method by Benson and Winawer, 2018 that takes measured retinotopy data as prior information to compute a quasi-conformal mapping from a retinotopy template to the subject anatomy (see ***Supplementary Figure 1***), which is implemented in the neuropythy package (0.12.5, https://pypi.org/project/neuropythy/). Resulting smooth retinotopy maps were then used to delineate V1 and estimate iso-eccentricity lengths (see Cortical distances along iso-eccentricity lines).

Surface reconstruction of the cerebral cortex was based on the MP2RAGE UNI image. The UNI image first underwent bias field correction using SPM12. The resulting image was fed into the recon-all pipeline in FreeSurfer (6.0.0, http://surfer.nmr.mgh.harvard.edu/) (Dale, Fischl, and Sereno, 1999; Fischl, Sereno, and Dale, 1999) with the hires flag to segment at the original voxel resolution (Zaretskaya et al., 2018). The brain mask during segmentation was computed from the second inversion image of the MP2RAGE acquisition using the SPM12 segmentation algorithm. The mask was defined by excluding all voxels exceeding the tissue class threshold of 10% in all non-white matter (WM) and non-gray matter (GM) tissue classes. The resulting GM boundary surfaces to WM and to cerebrospinal fluid (CSF; pial boundary surface) were manually corrected, and extra care was applied to correct the pial surface near the sagittal sinus. Final GM/WM surfaces were shifted 0.5 mm inward to counteract a potential segmentation bias using FreeSurfer with MP2RAGE (Fujimoto et al., 2014). Surface meshes were then smoothed using mris_smooth with 2 smoothing iterations implemented in FreeSurfer. Smoothed surfaces were upsampled to an average edge length of around 0.3 mm. Based on the registration to the functional time series without task (see further below), boundary surfaces were transformed to the space of the functional image by applying the deformation field to vertex coordinates using linear interpolation. To consider different distortions in anatomical and functional images, we aligned registered surface meshes with the GM/WM border found in the functional images (see ***Supplementary Figure 2***) using the GBB package (0.1.6, https://pypi.org/project/gbb/). Finally, nine meshes were positioned equidistantly between both boundary surfaces, resulting in 11 cortical layers for analysis.

For the registration of whole-brain anatomy and functional time series without task that was acquired in the same session, we first transformed the anatomical image to the functional space using the header information. Both images were then rigidly registered using ANTs (2.3.1, http://stnava.github.io/ANTs/). For preparation, the brain in both images were masked, and the functional data was bias field corrected (Tustison et al., 2010).

All other functional time series were registered to the functional time series without task in an equivalent way with the only difference that a nonlinear registration was applied using the Symmetric Normalization (SyN) algorithm (Avants et al., 2008) implemented in ANTs to account for different distortions between sessions. All runs were independently registered.

Based on the computed registrations, we deformed surface meshes using linear interpolation into the target space from individual functional runs to sample data onto single cortical layers using linear interpolation.

### MTF model fitting using MCMC

In our analysis, we assume that the complex-valued fMRI signal can be described as convolution with a point spread function kernel *PSF*_*BOLD*_(*x, t*) (Parkes et al., 2005) parameterized by cortical distance *x* and time *t*, which is the Fourier transform of the modulation transfer function *MT F*_*BOLD*_(*k, ω*) with spatial frequency *k* in cycles/mm and temporal frequency *ω* in rad/s. Our MTF model then describes the BOLD amplitude at stimulation frequency *c*_*s*_(*k*_*ecc*_) in dependence on the spatial frequency of the induced traveling wave *k*_*ecc*_, which is determined by cortical position and number of wedges used for stimulation. The expected value of the BOLD magnitude *E*[*c*_*s*_(*k*_*ecc*_)] is then given by assuming a Rice distribution (Gudbjartsson and Patz, 1995)

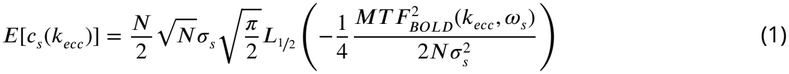

with baseline offset *σ*_*s*_ resulting from Gaussian white noise added to the complex-valued fMRI time series. A detailed derivation of the MTF model can be found in Appendix: Derivation of the modulation transfer function model. For *MT F*_*BOLD*_, we assume a Gaussian function with free parameters *β*_*BOLD*_ (amplitude of *MT F*_*BOLD*_), representing the BOLD response amplitude to a spatially homogeneous neuronal response (*k*_*ecc*_ = 0), and *σ*_*BOLD*_ (width of *MT F*_*BOLD*_). The effective resolution of the BOLD signal can then be described by the full width at half maximum (FWHM) of *PSF*_*BOLD*_(*x, t*) that relates to *σ*_*BOLD*_ by

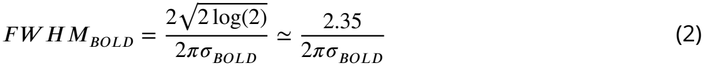

For model fitting, we used the BOLD amplitudes at stimulation frequency averaged across both sessions acquired with the same MR pulse sequence (see Percent signal changes across cortical depth) and estimated for each V1 location the corresponding spatial frequency of the traveling wave (see Cortical distances along iso-eccentricity lines). Furthermore, we only included data points from V1 that were inside the FOV of all GE- and SE-BOLD sessions. Selected amplitudes were binned (*n* = 200) based on their associated spatial frequencies into equidistant spatial frequency intervals.

We used Bayesian hierarchical modeling to estimate the parameters *β*_*BOLD*_, *σ*_*BOLD*_, and *σ*_*s*_ at the subject and population level. In Bayesian modeling, we estimate probability distributions for the parameters of interest (posterior distribution) by updating defined prior distributions with the model likelihood according to observed data. ***Figure 3*** shows the graph representation of the defined model. We used Gamma distributions *Ga*(*α, β*) with shape *α* and rate *β* parameters, respectively, as prior distributions at the subject level since all parameters can only have non-negative values.

**Figure 3.**
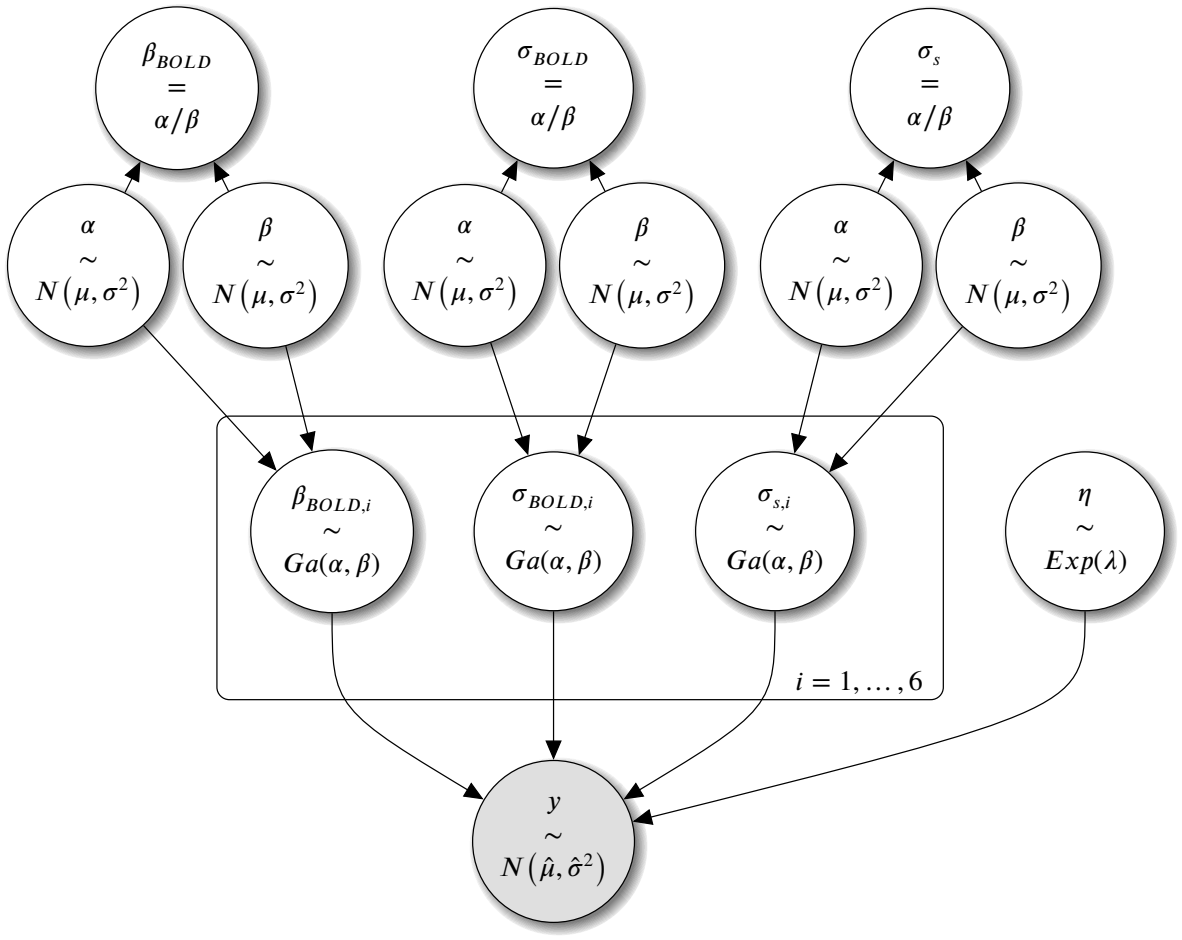
Graphical representation of the applied Bayesian hierarchical regression model. For each subject *i* = 1, …, 6, acquired data was modeled by the MTF model that is derived in Appendix: Derivation of the modulation transfer function model and has the free parameters *β*_*BOLD,i*_, *σ*_*BOLD,i*_ and *σ*_*s,i*_. We used Gamma distributions *Ga*(*α, β*) as prior distributions with shape parameter *α* > 0 and rate parameter *β* > 0 (subject level). *α* and *β* were drawn for each parameter from individual hyperprior distributions that were assumed to follow normal distributions (population level). In ***Table 1***, the settings for prior and hyperprior distributions are listed. We used ^*α*^∕_*β*_ as mean parameter estimates at the population level. Finally, the model likelihood *y* was computed assuming a normal distribution 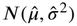 with the exponential distribution *Exp*(*λ*) with rate parameter *λ* > 0 being the prior distribution of the variance of the normal distribution.

In hierarchical modeling, we assume that probability distributions of parameters at the subject level can be derived from a common distribution at the population level. In our model, *α* and *β* were taken from normal distributions *N*(*µ, σ*^2^) that were used as hyperprior distributions at the population level. In ***Table 1***, the settings for prior and hyperprior distributions are listed. At the population level, the distribution mean of Gamma distributions is characterized by the ratio *α*∕*β*. For *σ*_*BOLD*_, the ratio was set to 0.16 cycles/mm, which corresponds to a FWHM of 2.34 mm that was taken from Shmuel et al., 2007.

**Table 1.**
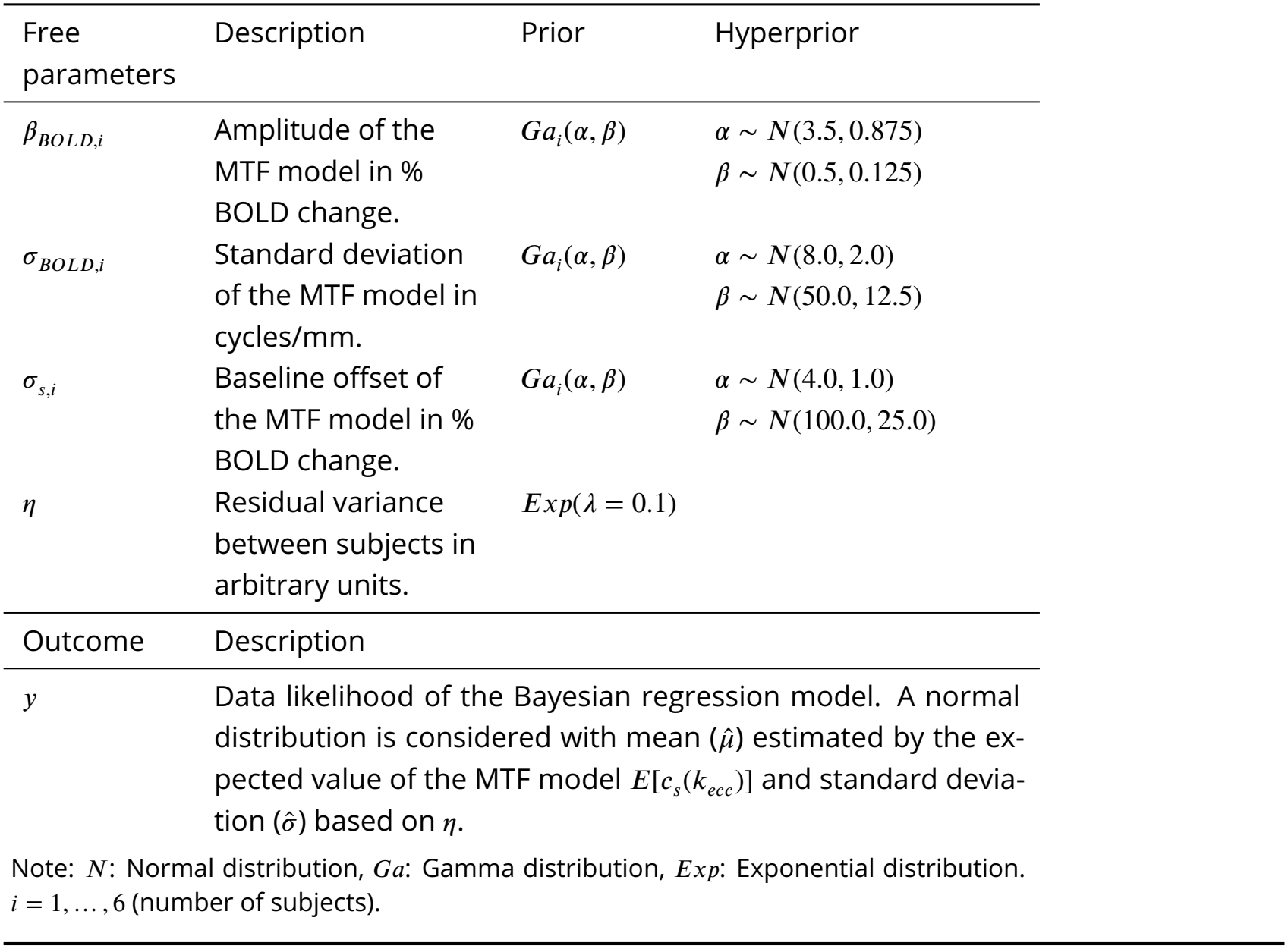
Model parameters. A graphical visualization of the model can be found in ***Figure 3***. More information about settings of prior and hyperprior distributions is stated in MTF model fitting using MCMC.

| Free parameters | Description | Prior | Hyperprior |
|---|---|---|---|
| *β*<sub>*BOLD,i*</sub> | Amplitude of the MTF model in % BOLD change. | *Ga*<sub>*i*</sub>(*α*, *β*) | *α* ∼ *N*(3.5, 0.875)<br>*β* ∼ *N*(0.5, 0.125) |
| *σ*<sub>*BOLD,i*</sub> | Standard deviation of the MTF model in cycles/mm. | *Ga*<sub>*i*</sub>(*α*, *β*) | *α* ∼ *N*(8.0, 2.0)<br>*β* ∼ *N*(50.0, 12.5) |
| *σ*<sub>*s,i*</sub> | Baseline offset of the MTF model in % BOLD change. | *Ga*<sub>*i*</sub>(*α*, *β*) | *α* ∼ *N*(4.0, 1.0)<br>*β* ∼ *N*(100.0, 25.0) |
| *η* | Residual variance between subjects in arbitrary units. | *Exp*(*λ* = 0.1) |  |
| Outcome | Description |  |  |
| *y* | Data likelihood of the Bayesian regression model. A normal distribution is considered with mean (*μ̂*) estimated by the ex-pected value of the MTF model *E*[*c*<sub>*s*</sub>(*k*<sub>*ecc*</sub>)] and standard devia-tion (*σ̂*) based on *η*. |  |  |

Note: *N*: Normal distribution, *Ga*: Gamma distribution, *Exp*: Exponential distribution. *i* = 1, … , 6 (number of subjects).

We assumed a normal distribution for the model likelihood, with the mean characterized by our MTF model and standard deviation determined by a noise distribution that accounts for unexplained variance at the population level.

For inference, we used MCMC sampling to directly sample from the joint posterior probability distribution and thus generate estimates of the marginal distributions for our parameters of interest. To ensure that parameter estimates accurately reflect the target distribution, MCMC samples must reach convergence (see MCMC diagnostics). For MCMC sampling, we used PyMC (4.2.0, https://pypi.org/project/pymc/) with the NUTS sampler (Hoffman and Gelman, 2014). We ran the sampler for 36,000 iterations and discarded the first 30,000 samples (burn-in period). The remaining 6,000 samples were thinned by only taking every third sample, which resulted in 2,000 samples for analysis.

## Results

### GE- and SE-BOLD maps

***Figure 4*** shows the amplitude and phase of Fourier components at stimulation frequency from runs with 2 and 14 wedges of GE- and SE-BOLD sessions for one participant (see ***Supplementary Figure 3*** for illustrations of all single participants). The phase responses in ***Figures 4A–D*** illustrate the expected traveling wave pattern with the direction of propagation from one vertical meridian to the other. As expected, traveling waves from runs with 14 wedges showed an increased spatial frequency. Note, however, that the local spatial frequency of traveling waves on the cortical surface also depends on the spatially varying cortical traveling distance in V1 in addition to the number of wedges (see Appendix: Derivation of the modulation transfer function model).

**Figure 4.**
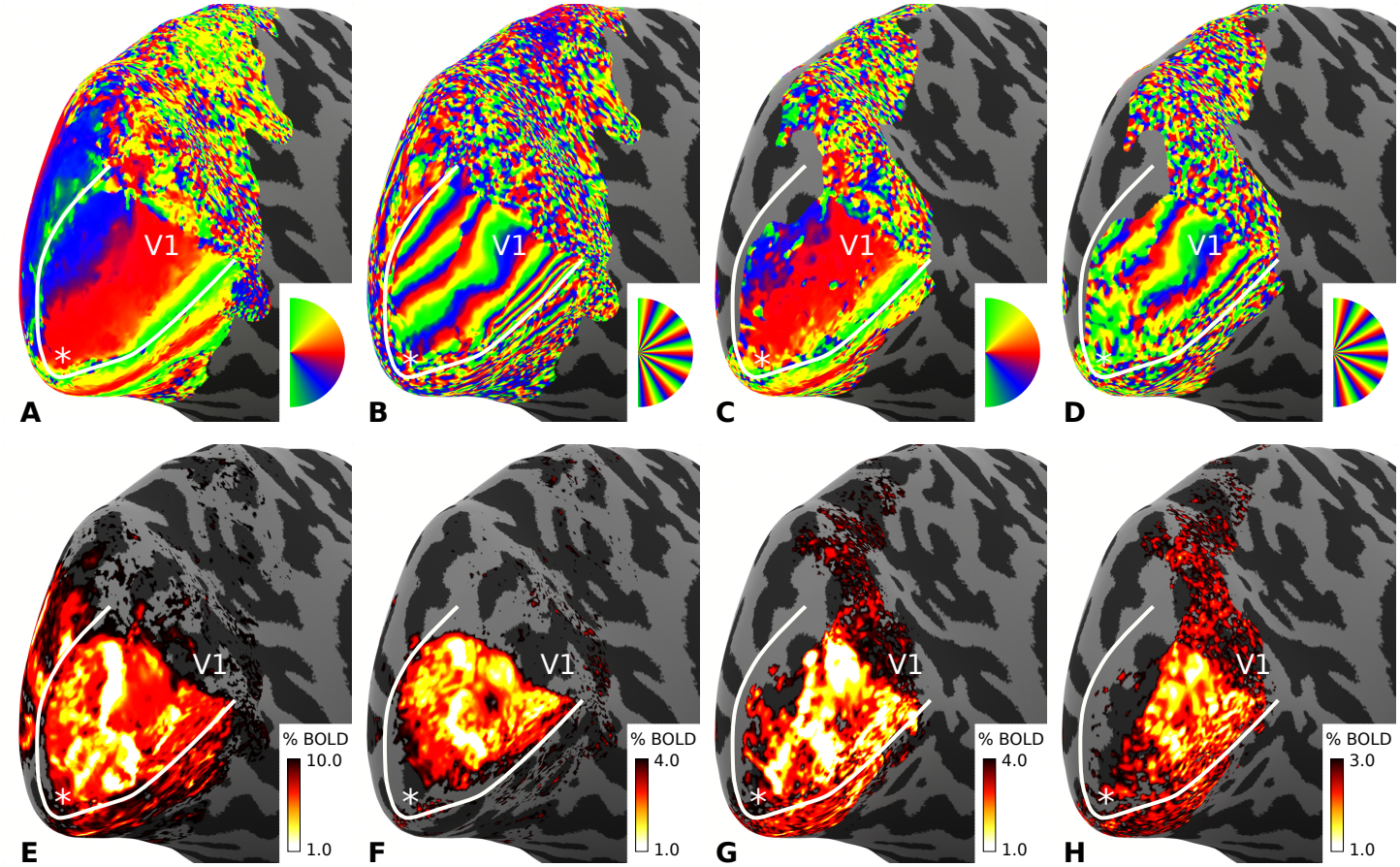
Exemplary phase responses and BOLD signal changes. For one representative participant (subject 6; left hemisphere), the phase of the BOLD signal at different spatial stimulation frequencies is shown on an inflated cortical surface mesh in **A–D. A** and **B** illustrate responses from GE-BOLD measurements for runs with 2 and 14 wedges, respectively. **C** and **D** show corresponding responses from SE-BOLD measurements. Associated BOLD magnitudes at stimulation frequency are shown in **E–H**. Note the different color scales used for magnitude data. The data was sampled at mid-cortical depth and averaged across two sessions. Solid white lines indicate the V1/V2 border that was based on a separate retinotopy session (see ***Supplementary Figure 1***). White asterisks mark the location of the foveal representation. Note that we acquired fewer slices for SE-BOLD measurements, which resulted in less coverage of V1 (e.g., see the area around the lower vertical meridian that was not covered in SE-BOLD measurements). Phase and magnitude responses from all single participants can be found in ***Supplementary Figure 3***.

For the runs with 14 wedges, no consistent phase responses were seen outside of V1. This absence in extrastriate cortical areas is most probably a consequence of the larger receptive field sizes outside of V1 (Dumoulin and Wandell, 2008). Note that we restricted our analysis to area V1. While the phase responses from SE-BOLD sessions were noisier (e.g., more phase wraps can be identified), a high degree of similarity between GE- and SE-BOLD phase maps can qualitatively be seen.

The amplitudes shown in ***Figures 4E–H*** demonstrate the higher signal changes for GE-in comparison to SE-BOLD sessions (see the different *y*-axis scalings). In addition, signal changes got smaller with increasing number of wedges, which is the expected behavior described in our MTF model. However, locations around the foveal representation showed an even larger dampening of signal changes with increasing number of wedges. This consistent finding across participants might reflect the further spatial frequency increase of traveling waves due to the smaller cortical distances toward the foveal representation. However, note that eye movements will have more impact on signal changes around the foveal representation due to differences in cortical magnification, which is not accounted for in the present study. Missing signal changes around the upper vertical meridian in SE-BOLD maps only resulted from the smaller FOV that was imaged due to SAR constraints (see Imaging).

While we depict the phase maps in ***Figure 4*** to illustrate the pattern of traveling waves, we only used the amplitude data for the MTF analysis.

### Percent signal changes across cortical depth

***Figures 5A–B*** show the amplitude at stimulation frequency as a function of cortical depth from GE- and SE-BOLD sessions, respectively. Both plots depict the mean amplitude averaged across all participants from individual runs with varying numbers of wedges. As before, GE-BOLD overall showed larger signal changes for all runs across participants. Furthermore, we also observed a steady decrease in signal changes with increasing number of wedges, which is the behavior that the MTF model exploits. In addition, we observed the typical increase of BOLD signal changes toward the pial surface, commonly attributed to contributions from macrovascular draining veins (Turner, 2002; Polimeni et al., 2010).

**Figure 5.**
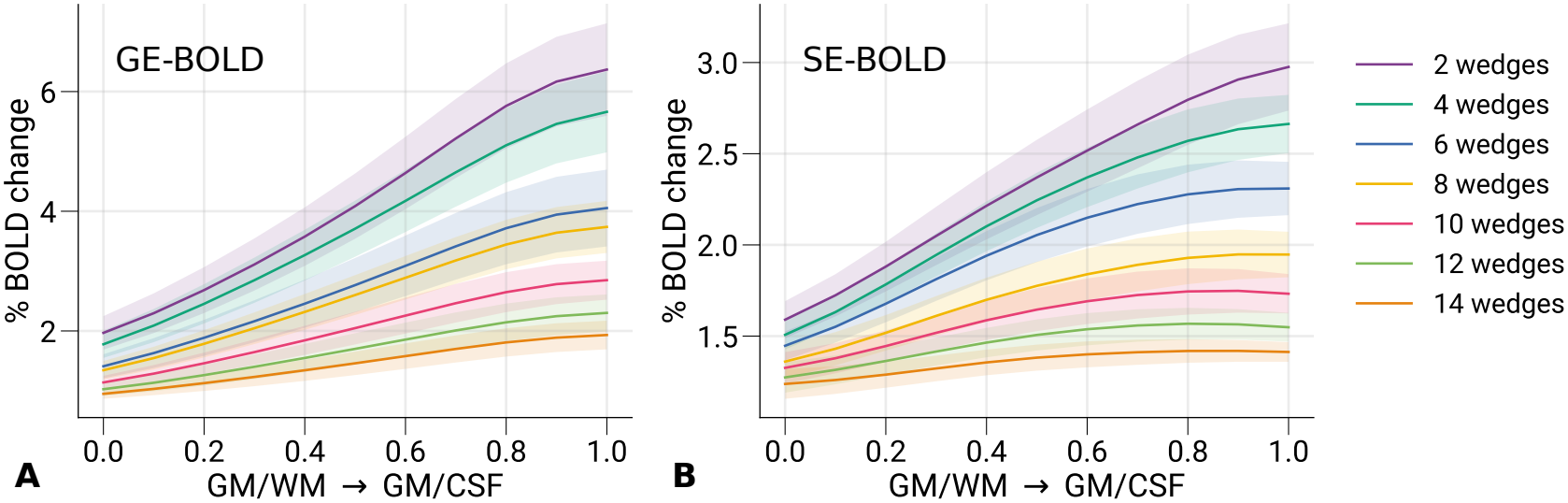
Cortical profiles of % BOLD signal changes. Mean signal changes across participants are shown for **A** GE- and **B** SE-BOLD sessions at different cortical depths. The typical increase of signal changes toward the pial surface can be seen, while the overall response diminishes with greater number of wedges, i.e., with higher spatial frequency of stimuli. Shaded areas denote the standard error of the mean across participants.

### Cortical distances along iso-eccentricity lines

Placing the MTF model in the context of cortical functional neuroanatomy, we aim to express the spatial frequency of traveling waves in terms of cycles per mm of cortical tissue. We, therefore, need an estimate of distances along the cortical surface for the paths of traveling waves. The induced traveling waves will travel in polar angle direction along a path with constant eccentricity (iso-eccentricity lines). We used polar angle and eccentricity maps from a separate retinotopy measurement for the estimation of distances. However, since eccentricity and polar angle maps are usually based on voxel-wise estimates, they are prone to noise in each, leading to discontinuous maps, which affects accurate length estimation. Therefore, we used the method by Benson and Winawer, 2018, which yielded smooth retinotopic maps with preserved topology. For each voxel in V1, we then estimated the geodesic path that connects both upper and lower vertical meridians of V1 while holding eccentricity constant. This analysis was done separately for each cortical depth.

In ***Figure 6***, a histogram for the length of iso-eccentricity lines (distance from one vertical meridian to the other) pooled across participants is shown that was estimated at mid-cortical depth. Since these lengths were calculated for each hemisphere separately, they correspond to the traveling wave propagation of one half stimulus cycle. The modes of the distribution of iso-eccentricity lengths for single participants are shown as colored vertical lines, demonstrating the variability of V1 size between participants.

**Figure 6.**
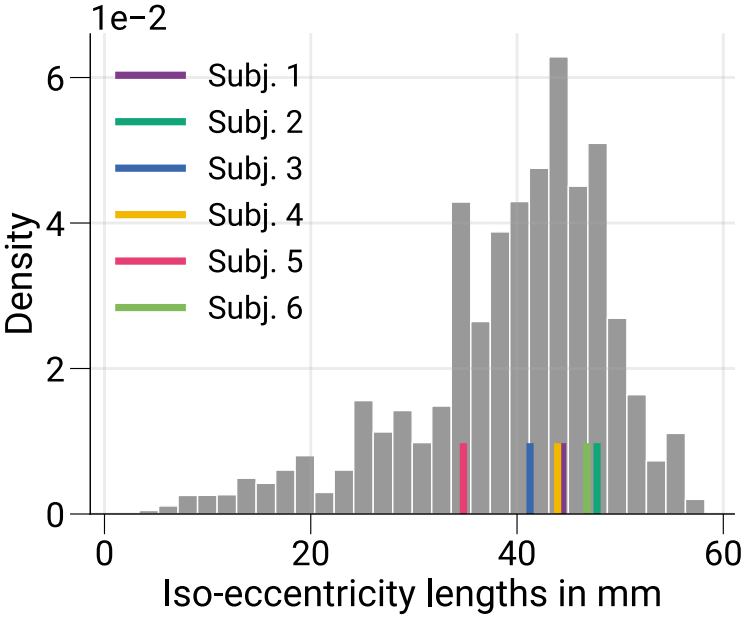
Lengths of iso-eccentricity lines in V1. The histogram shows the normalized distribution of path lengths for curves on the cortical surface that run between vertical meridians, i.e., V1/V2 borders, within V1 at constant eccentricity (iso-eccentricity lengths), which is the direction of wave propagation of induced traveling waves. Lengths are shown for surface coordinates at mid-cortical depth. Data were pooled across participants, and only the stimulated part of V1 was considered. Colored vertical lines indicate the distribution modes for single participants.

### MCMC diagnostics

We used the amplitudes at stimulation frequency (see Percent signal changes across cortical depth and corresponding spatial frequencies (see Cortical distances along iso-eccentricity lines) for fitting our MTF model using a hierarchical Bayesian regression model. The posterior probability distributions for the parameters of interest were estimated using MCMC sampling. To inspect that the sampled distributions approximate the target distributions, we must ensure the parameter samples converge, i.e., traces of parameter samples are stationary.

***Figure 7*** shows exemplary diagnostics for parameter traces at the population level estimated at mid-cortical depth to demonstrate the degree of reached convergence. The left column directly shows sampled parameter traces that qualitatively illustrate the absence of slow drifts (see ***Supplementary Figure 4*** for parameter traces at the subject level). The next column shows the auto-correlation of these traces, which illustrates that already after a few lags, the autocorrelation is indistinguishable from that of a white noise process. We also computed the Geweke diagnostics, a *z*-test that compares the first 10% to the last 50% of parameter traces (Geweke, 1991). In the last column, we show scatter plots that qualitatively demonstrate no covariance between parameter estimates.

**Figure 7.**
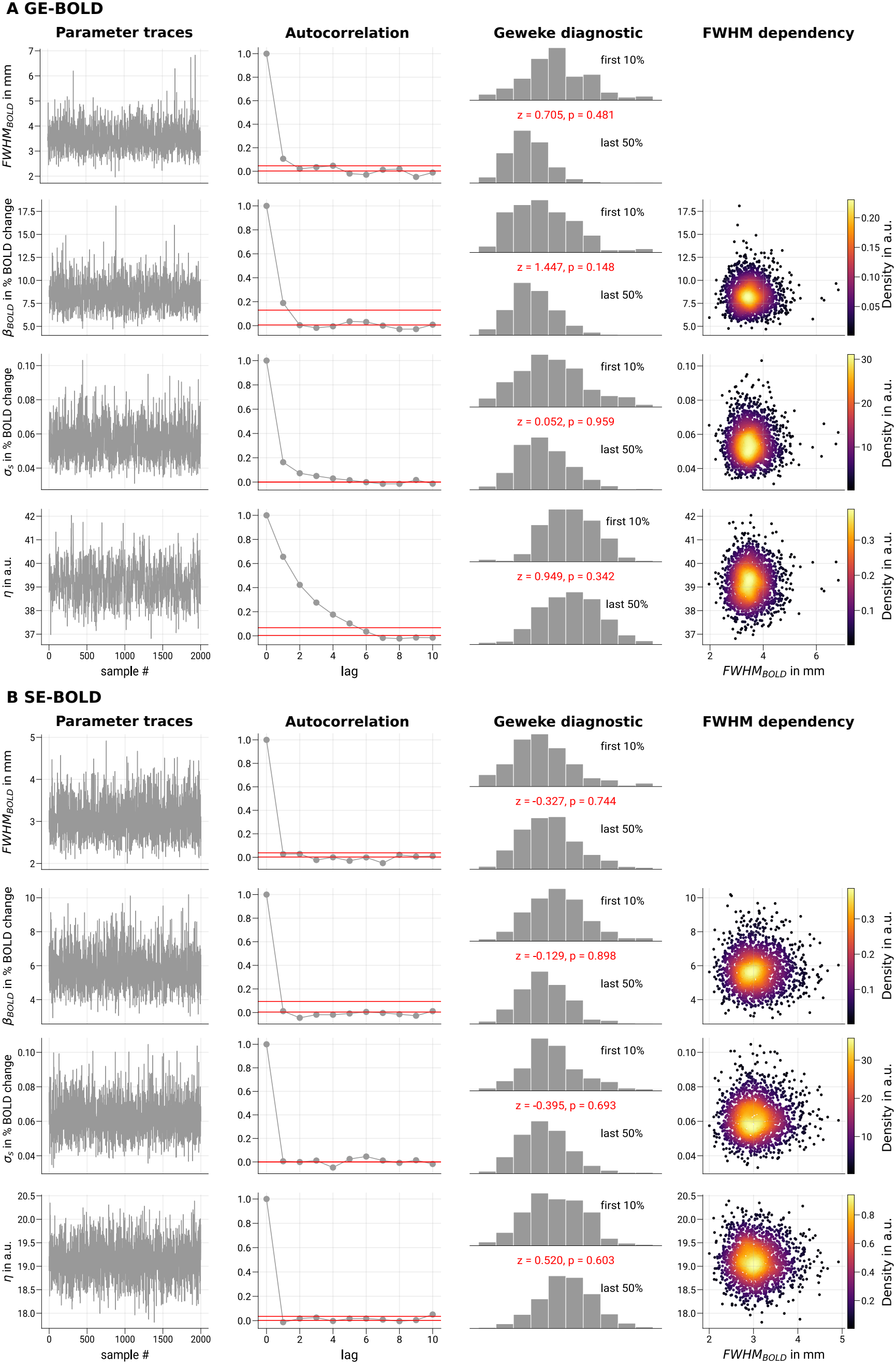
Convergence diagnostics of Markov chain Monte Carlo sampling. For Markov chain Monte Carlo (MCMC) sampling, enough sampling iterations must be performed to ensure convergence that allows an unbiased sampling from the posterior distribution. Indications for convergence are stationarity of sampled parameters and a rapidly decreasing autocorrelation with increasing lag relative to the total number of samples. Exemplary diagnostics are shown for GE- and SE-BOLD in **A** and **B**, respectively, for data sampled at mid-cortical depth. The first column shows traces of sampled parameter estimates at the population level (see MTF model fitting using MCMC for further details of the Bayesian regression model). Corresponding trace plots for single participants are shown in ***Supplementary Figure 4***. No trends are visually noticeable, which points toward reached stationarity. The second column shows sample autocorrelations. Horizontal red lines indicate the 95% confidence interval around 0 for a white noise process, which shows the bounds of expected autocorrelation found in uncorrelated white noise. The third column shows the Geweke convergence diagnostic, a *z*-test (*z*-scores and corresponding *p*-values shown in red) that tests whether the means of the first 10% and last 50% of samples are different. The last column shows sample covariations to the parameter of interest *F W HM*_*BOLD*_.

### Estimated MTF parameters

For each cortical depth independently, we estimated posterior probability distributions for our parameters of interest using MCMC sampling. ***Figure 8*** shows the distributions of parameter estimates at the population level for *β*_*BOLD*_, *F W HM*_*BOLD*_ and *σ*_*s*_ as detailed in ***Table 1*** from GE- and SE-BOLD sessions estimated at mid-cortical depth as scatter plots for comparison (see ***Supplementary Figure 5*** for an illustration of parameter fits at the subject level). In ***Figure 8A***, it can be seen that most amplitudes of *MT F*_*BOLD*_ were larger for GE-BOLD, which reflects the BOLD signal changes at stimulation frequency illustrated in GE- and SE-BOLD maps and Percent signal changes across cortical depth, respectively.

**Figure 8.**
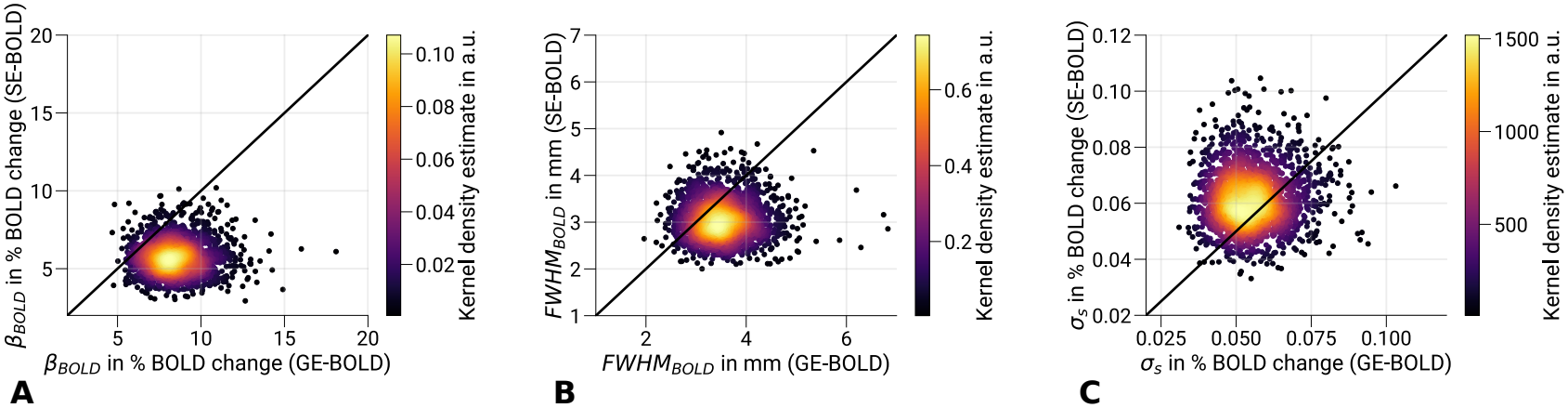
Comparison of parameter estimates. Scatter plots show the comparisons between GE- and SE-BOLD parameter estimates for **A** *β*_*BOLD*_, **B** *F W HM*_*BOLD*_ and **C** *σ*_*s*_ at the population level. Data were sampled at mid-cortical depth. Black lines show the identity line. The majority of individual *β*_*BOLD*_ and *F W HM*_*BOLD*_ samples are larger for GE-than for SE-BOLD shown in **A** and **B**, respectively. In **C**, samples show slightly larger values for SE-compared to GE-BOLD. Mean estimates for GE- and SE-BOLD are stated in ***Table 2***. A comparison of parameter estimates at the subject level is shown in ***Supplementary Figure 6***.

Similar results were found for *F W HM*_*BOLD*_ shown in ***Figure 8B***, which is the used metric to characterize the spatial specificity of the functional signal. It illustrates that the majority of individual samples showed a wider point spread function width for GE-than for SE-BOLD. However, the distributions also covered a wide range of widths. In ***Supplementary Figure 6***, the comparison of parameter distributions between GE- and SE-BOLD is shown at the subject level, demonstrating a high variability between participants. But still, all participants but one revealed a wider *PSF*_*BOLD*_ width for GE-BOLD at the subject level. Interestingly, the participant who showed the opposite effect also had the smallest difference in *MT F*_*BOLD*_ amplitudes between GE-BOLD and SE-BOLD, which might point toward poor data quality (see Discussion for an extended discussion about data quality). Note that being more robust against potential outliers at the subject level was one of the main reasons to use a hierarchical modeling approach. The baseline noise level shown in ***Figure 8C*** did not show conclusive differences between GE- and SE-BOLD at the population and subject level (see ***Supplementary Figure 6***).

**Table 2.** The median of parameter estimates at different cortical depths and their dependency on static susceptibility effects. For GE- and SE-BOLD, median parameter estimates (*β*_*BOLD*_, *F W HM*_*BOLD*_, *σ*_*s*_) at the population level are listed for three different cortical depths (0%: GM/CSF boundary, 50%: mid-cortical depth, 100%: GM/WM boundary). Furthermore, parameter estimates from cortical locations with high (hypointense EPI) and low (hyperintense EPI) static susceptibility effects and, therefore, much or little contributions from large veins are shown. For each participant, these locations were defined by taking only data in V1 that belong to the lower and upper 5th percentile of a bias-corrected 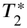-weighted temporal mean from a separately acquired EPI time series (see ***Figure 10***). It is known that contributions from large veins manifest themselves as dark intensities in 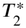-weighted images due to short blood 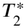 at high magnetic fields (Yacoub et al., 2001) and increased susceptibility differences to surrounding tissue (Kay et al., 2019). GM: gray matter, WM: white matter, CSF: cerebrospinal fluid.

|  |  | *β*<sub>*BOLD*</sub><br>in % BOLD change | *FWHM*<sub>*BOLD*</sub><br>in mm | *σ*<sub>*s*</sub><br>in % BOLD change |
|---|---|---|---|---|
| GE-BOLD | Cort. Depth 0% | 11.99 [8.64; 17.19] | 3.98 [2.99; 5.55] | 0.08 [0.06; 0.11] |
|  | Cort. Depth 50% | 8.35 [5.90; 11.62] | 3.49 [2.64; 4.64] | 0.05 [0.04; 0.08] |
|  | Cort. Depth 100% | 4.96 [3.35; 7.15] | 3.27 [2.50; 4.43] | 0.05 [0.03; 0.06] |
|  | hypointense EPI | 9.02 [6.50; 12.68] | 3.79 [2.85; 5.23] | 0.07 [0.05; 0.10] |
|  | hyperintense EPI | 8.14 [5.84; 11.58] | 3.48 [2.63; 4.80] | 0.05 [0.04; 0.08] |
| SE-BOLD | Cort. Depth 0% | 6.62 [4.69; 9.38] | 3.78 [2.85; 5.16] | 0.07 [0.05; 0.10] |
|  | Cort. Depth 50% | 5.68 [3.90; 8.23] | 3.03 [2.34; 4.06] | 0.06 [0.04; 0.08] |
|  | Cort. Depth 100% | 3.98 [2.60; 5.75] | 2.95 [2.26; 3.88] | 0.06 [0.04; 0.09] |
|  | hypointense EPI | 5.94 [4.15; 8.43] | 3.64 [2.74; 4.94] | 0.07 [0.05; 0.10] |
|  | hyperintense EPI | 5.86 [3.98; 8.25] | 3.15 [2.39; 4.24] | 0.05 [0.04; 0.08] |

Note: Numbers in square brackets denote the 95% credible interval of parameter estimates.

In ***Figure 9***, the median of estimated parameter distributions with associated 95% credible interval is shown across cortical depth. We used the median to be more robust against potentially remaining non-stationarities in parameter traces. The *MT F*_*BOLD*_ amplitudes shown in ***Figure 9A*** reflect the increase of BOLD signal changes toward the pial surface shown in Percent signal changes across cortical depth. ***Figure 9B*** also shows an increase toward the pial surface for the *PSF*_*BOLD*_ width. The large variability between participants is reflected in the largely overlapping credible intervals. However, SE-BOLD showed a small but consistently smaller *PSF*_*BOLD*_ width across the cortical ribbon. The difference of parameter medians for the *PSF*_*BOLD*_ width at mid-cortical depth was 0.46 mm, which is larger but comparable to the difference between GE- and SE-BOLD stated in Chaimow et al., 2018. Note that voxels with macrovascular contributions were excluded in their comparison, which was not done in the current main analysis.

**Figure 9.**
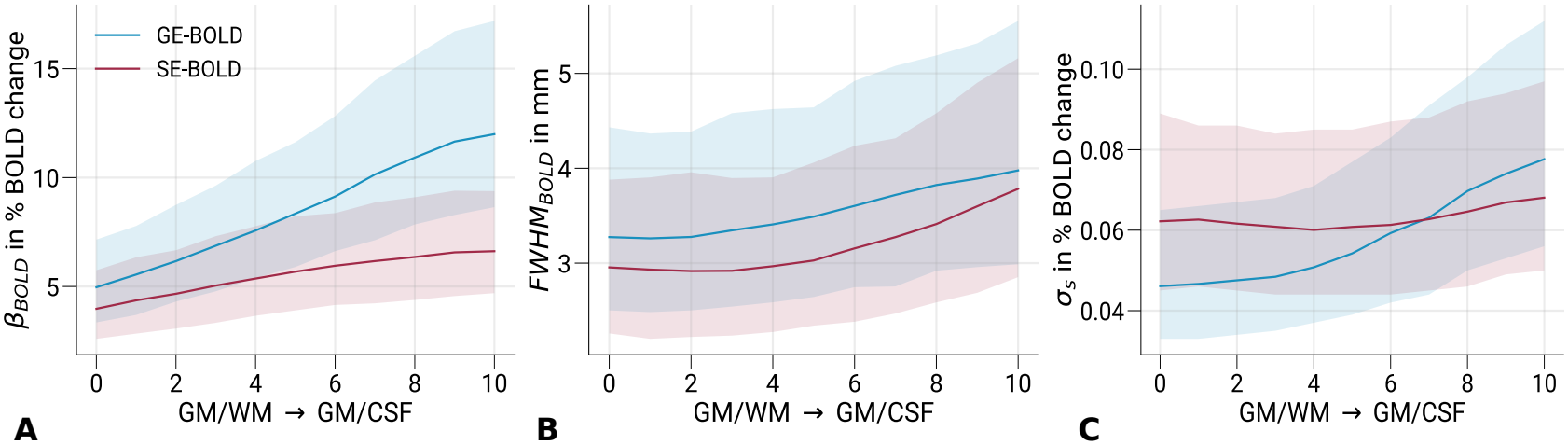
Cortical profiles of parameter estimates. Parameter estimates of **A** *β*_*BOLD*_, **B** *F W HM*_*BOLD*_ and **C** *σ*_*s*_ are shown at different cortical depths. In **A**, the expected increase of signal amplitudes toward the pial surface can be seen. In **B**, both GE- and SE-BOLD show an increase of the *F W HM*_*BOLD*_ toward the pial surface, while SE-BOLD is marginally more specific at all cortical depths. In **C**, the GE-BOLD baseline offset *σ*_*s*_ shows a larger but non-significant change across cortical depth. Single lines show the median parameter estimate at the population level with associated 95% credible interval. Corresponding values at three cortical depths are listed in ***Table 2***.

Interestingly, the difference was also smaller than the difference across cortical depth, which was 0.71 mm and 0.83 mm for GE- and SE-BOLD, respectively. A summary of mean parameter estimates can be found in ***Table 2***.

### MTF in proximity to veins

In the preceding analysis, we used all available data in V1. However, the vascular structure is known to be heterogeneous both in radial, i.e. across cortical depth, and tangential, i.e. along the cortical surface, directions. Here, we added an analysis comparing the MTF properties from V1 locations depending on their venous contributions. The analysis was restricted to data sampled at mid-cortical depth to avoid introducing biased estimates of venous contribution due to partial voluming with WM and CSF.

We leveraged the fact that deoxygenated blood in venous compartments exhibits a short 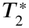 at high magnetic fields (Yacoub et al., 2001), resulting in signal loss within and around veins due to pronounced susceptibility differences relative to gray matter tissue. Consequently, veins can be identified as hypointense regions in 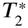-weighted images (Ogawa and Lee, 1990; Shmuel et al., 2010; Kay et al., 2019). We used the 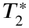-weighted EPI time series without task acquired in the first session (TR = 3000 ms, number of volumes = 200), which has the additional advantage that spatial distortions due to the low bandwidth in phase-encoding direction are matched to other functional measurements. The motion-corrected time series was highpass filtered (cutoff frequency: 1/270 Hz), and the temporal mean was calculated. The mean image was further bias field corrected (Tustison et al., 2010) and sampled at mid-cortical depth.

***Figure 10*** shows the bias field corrected temporal mean for one typical participant (same participant as in ***Figure 4***). A variable intensity pattern can be seen in area V1. The green arrow points to a location putatively dominated by macrovascular contributions. As expected, this corresponds to locations with the greatest BOLD signal changes (Olman, Inati, and Heeger, 2007), as shown in ***Figure 4***.

**Figure 10.**
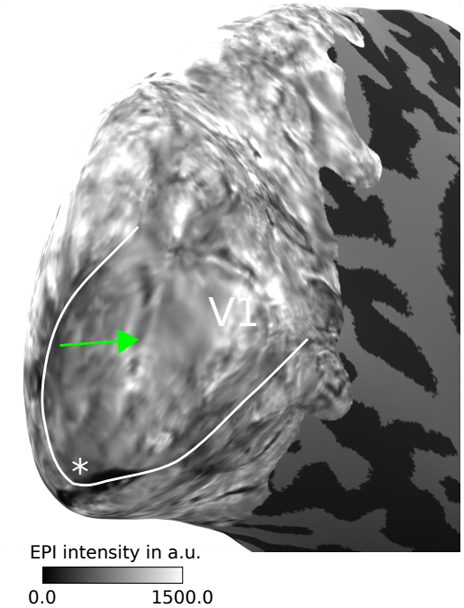
Exemplary visualization of static susceptibility maps. For one participant (subject 6), the bias-field corrected temporal mean of the EPI scan without task is shown on the inflated surface. Data were sampled at mid-cortical depth. It is known that contributions from large veins manifest themselves as dark intensities in 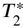-weighted data due to short blood 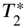 at high magnetic fields (Yacoub et al., 2001) and increased susceptibility differences between veins and surrounding tissue (Kay et al., 2019). The green arrow points to a putative large vein visible as a dark line. Compared to ***Figure 4***, this location corresponds to the area where BOLD signal changes peak in V1 as expected from voxels near large veins (Olman, Inati, and Heeger, 2007). The solid white line indicates the V1/V2 border that was based on a separate retinotopy session (see ***Supplementary Figure 1***). The white asterisk marks the location of the foveal representation.

We defined regions with little and significant vein contributions by finding the areas of V1 that belong to the upper and lower 5th percentile of the temporal mean, respectively. MTF analyses were then performed independently for data in both regions.

***Figure 11*** shows that both GE- and SE-BOLD are affected by macrovascular contributions. *MT F*_*BOLD*_ amplitudes and *PSF*_*BOLD*_ widths increased for locations with venous contributions; see also ***Table 2***. Interestingly, the baseline noise level *σ*_*s*_ also increased in regions with venous contributions, which might be explained by physiological noise contributions induced by cardiac pulsation that are localized around large arteries and draining veins (Glover, Li, and Ress, 2000; Caballero-Gaudes and Reynolds, 2017).

**Figure 11.**
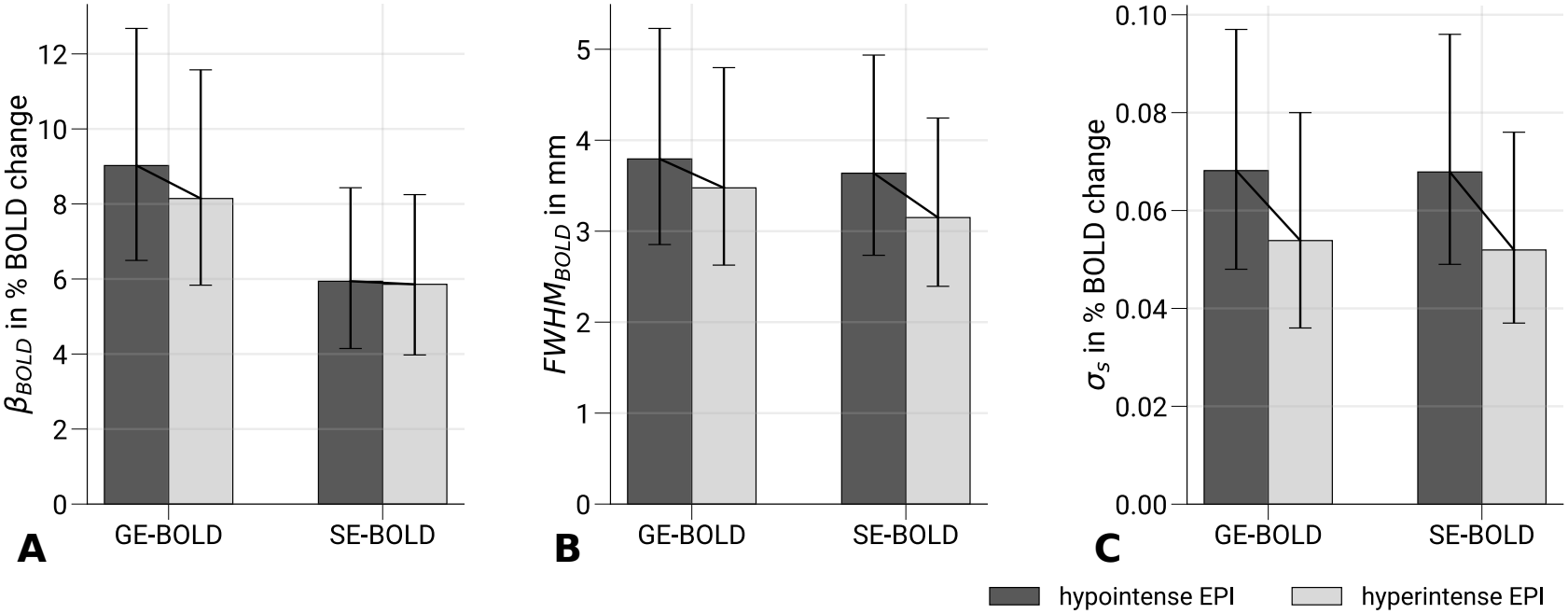
Dependency of parameter estimates to static susceptibility effects. Bar graphs show the comparison of **A** *β*_*BOLD*_, **B** *F W HM*_*BOLD*_ and **C** *σ*_*s*_ parameter estimates of GE- and SE-BOLD, respectively, between areas with high (hypointense EPI) and low (hyperintense EPI) static susceptibility effects, and therefore, much or little contributions from large veins. Static susceptibility effects were estimated from the temporal mean of separately acquired 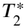-weighted EPI time series where locations with high static susceptibility manifest themselves as dark intensities (see ***Figure 10***). More precisely, hypointense and hyperintense EPI areas, respectively, were defined by taking only data in V1 that belong to the lower and upper 5th percentile of the temporal mean. Single bars show the median parameter estimate at the population level with associated 95% credible interval. Corresponding values are listed in ***Table 2***. It can be seen that locations with more contributions from large veins are less specific **(B)** and overall show higher noise levels **(C)**.

## Discussion

We estimated and compared the spatial specificity of GE- and SE-BOLD at 7 T at different cortical depths in the direction parallel to the cortical surface. At mid-cortical depth, we measured PSF widths of 3.49 mm and 3.03 mm for GE- and SE-BOLD (see ***Table 2***), respectively. Across cortical depth, the PSF width of SE-BOLD is smaller compared to GE-BOLD. However, both acquisition techniques show larger PSF widths toward the pial surface (see ***Figure 9***). The expected higher specificity of SE-BOLD originates from the features that intravascular signal contributions become negligible at higher fields due to the shortening of blood *T*_2_ (Thulborn et al., 1982; Yacoub et al., 2001) and that extravascular signal contributions from around larger veins are refocused (Boxerman et al., 1995; Uludağ, Müller-Bierl, and Uğurbil, 2009). The increase of the PSF for both GE- and SE-BOLD shows that macrovascular draining vein contributions, which lead to a spatial spread of the GE-BOLD signal, also affects SE-BOLD. This might be either introduced by remaining intravascular signal contributions or by additional 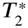-weighting along the EPI readout around the spin echo. Broadened PSFs close to veins at mid-cortical depth support this interpretation (see ***Figure 11***).

Our results therefore show that the spatial specificity of BOLD acquisitions generally depend on the proximity to local macrovasculature, which is in line with the usual recommendation of leaving out voxels in the analysis with venous contributions (Olman, Inati, and Heeger, 2007) and sampling data far away from the pial vasculature (Nasr, Polimeni, and Tootell, 2016) to increase the effective spatial specificity of the BOLD signal.

The PSF of the BOLD response was already estimated in different studies at different magnetic field strengths (Engel, Glover, and Wandell, 1997; Olman, Moortele, and Uğurbil, 2004; Parkes et al., 2005; Shmuel et al., 2007; Panchuelo et al., 2015; Cheng, 2016; Chaimow et al., 2018; Vizioli et al., 2020; Fracasso, Dumoulin, and Petridou, 2021). While the majority of studies focused on the GE-BOLD signal, a subset also included a comparison with SE-BOLD. All studies have in common that they targeted the visual cortex either exploiting its retinotopic organization (phase-encoded stimuli, grid-like stimuli, point-like stimuli of varying sizes, edge responses) or its known columnar organization (ocular dominance columns in V1, stripes in V2). However, exact methods, nominal voxel resolutions, and handling of venous contributions varied between studies, limiting their comparability.

In our study, we employed a phase-encoded paradigm that was similarly used by Engel, Glover, and Wandell, 1997 at 1.5 T, Parkes et al., 2005 at 3 T and Olman, Moortele, and Uğurbil, 2004 at 7 T, which all revealed a PSF width of around 3–4 mm that was also confirmed by our study. Among these studies, our methodology is closest to Parkes et al., 2005. However, our analysis differs notably in two aspects.

First, the absolute value of the PSF width depends on the accurate estimation of distances on the cortical surface. Parkes et al., 2005 estimated iso-eccentricity lengths based on Euclidean distances, which might be biased due to the high degree of cortical convolution in V1. In contrast, we based our estimates on geodesic distances, which revealed approximately twice as long distances. However, this only affects the absolute values of PSF width but not the comparison between GE- and SE-BOLD. Second, our MTF model (see Appendix: Derivation of the modulation transfer function model) included the estimation of non-zero baseline levels that can differ between GE- and SE-BOLD due to different noise contributions.

Parkes et al., 2005 also included a comparison between sequences and revealed PSF widths of 3.9 mm and 3.4 mm for GE- and SE-BOLD at 3 T, respectively. Interestingly, our study found a similar small difference in PSF width of around 0.5 mm. In a study by Chaimow et al., 2018, the PSF width of GE- and SE-BOLD was compared at 7 T exploiting the organization of ocular dominance columns in V1 and excluding voxels with macrovascular contributions. This study confirmed the only small PSF width decrease for SE-BOLD compared to GE-BOLD.

Recent studies predicted spatiotemporal BOLD responses using a physiologically-based model of the hemodynamic response (Aquino et al., 2012). A cortical depth-dependent analysis of the BOLD pattern found a linear increase of the spatial spread across cortical depth toward the pial surface (Puckett et al., 2016; Lacy et al., 2022). Another recent study by Fracasso, Dumoulin, and Petridou, 2021 analyzed the GE-BOLD PSF across cortical depth, exploiting the columnar organization of V2. They also showed a linear increase of PSF width toward the pial surface by about 1 mm with absolute values in the range of 0.8–1.8 mm. We confirm an increase of PSF width toward the pial surface in our data with the same order of magnitude. Furthermore, we show the first estimate of PSF widths for SE-BOLD across cortical depth and demonstrate a similar increase across cortical depth. Nevertheless, absolute values are less comparable due to the large differences in the experimental design, e.g., a differential paradigm that substracts responses to orthogonal conditions was used in Fracasso, Dumoulin, and Petridou, 2021, which is expected to already suppresses signal contributions not specific to the experimental condition.

Due to the use of single-shot echo-planar imaging (see Imaging), we underestimated the spatial specificity of SE-BOLD compared to a signal with pure *T*_2_-weighting. We aimed to perform an analysis at different cortical depths with high spatial resolution. Therefore, we used a long EPI readout that significantly affects the 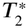 contribution of the signal (Keilholz et al., 2005; Goense and Logothetis, 2006; Norris, 2012; Van Horen et al., 2023). Furthermore, to have an acceptable signal-to-noise ratio (SNR), we had to decrease the echo time below the time for optimal BOLD contrast (Yacoub et al., 2003), which will also increase intravascular signal contributions. We note that 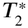 contamination during the readout is a general problem for high-resolution EPI-based SE-BOLD measurements. A discussion of the signal and noise characteristics of the outer *k*-space lines is, however, beyond the scope of the current study.

The cortical profiles of SE-BOLD signal changes (see ***Figure 5***) directly illustrate these limitations. Ideally, acquisitions dominated by microvascular contributions would exhibit a peak in signal changes in the middle cortical layers, reflecting the higher vascular density in layer 4 (Duvernoy, Delon, and Vannson, 1981; Weber et al., 2008). However, such a peak has been reported primarily in animal studies (Goense and Logothetis, 2006; Harel et al., 2006). Crucially, these studies used short or segmented readouts to minimize 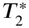-weighting around the spin echo. In contrast, our group-averaged SE-BOLD signal profiles increased toward the pial surface, suggesting substantial residual macrovascular contributions, potentially arising from 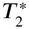-weighting during the readout and persistent intravascular signal.

Additional limitations are specific to the SE-BOLD acquisition. First, SAR constraints restricted the number of slices. Since inflow effects are most pronounced in the outer slices of the imaging slab, their impact likely differs between GE- and SE-BOLD. Second, off-resonance effects can have a stronger impact on SE-BOLD because of the lower bandwidth of the refocusing pulse, which may lead to a mismatch between excitation and refocusing slice profiles. We attempted to mitigate these effects by using an interleaved slice acquisition order.

We note that these limitations are inherent to high-resolution SE-BOLD imaging with single-shot EPI. Our results should therefore be understood as characterizing the SE-BOLD signal that is achievable in practice rather than an idealized *T*_2_-weighted spin-echo contrast.

Regarding these limitations, there are other alternatives for *T*_2_-weighted imaging at high magnetic field strengths. One possibility would be to use a 3D EPI with multiple refocusing pulses in partition direction, called 3D-GRASE (Oshio and Feinberg, 1991; Beckett et al., 2020). This readout scheme allows the selection of an inner volume (Feinberg et al., 1985), which can be used to reduce the in-plane echo train length. However, introducing multiple refocusing pulses will induce stimulated echoes and therefore additional *T*_1_-weighting across spatial frequencies, making the overall contrast more complex (Kemper et al., 2015; Kemper et al., 2016; Scheffler, Engelmann, and Heule, 2021). Alternatively, the 2D EPI readout can be segmented (Han et al., 2021). While this would be a straightforward approach to reduce the echo train length, the readout will be more susceptible to physiological noise, which introduces phase variations between shots and manifests itself as ghosting artifacts. Furthermore, image segmentation will lead to less time-efficient sampling and increased SAR for the same imaging volume. Other less SAR-intensive approaches that show a *T*_2_-weighted contrast are steady-state free precession (SSFP) or *T*_2_-prepared GE-BOLD, which have their own shortcomings and are further discussed in (Goa et al., 2014; Scheffler and Ehses, 2016; Scheffler et al., 2019; Ramadan et al., 2025; Pfaffenrot et al., 2021). In this regard, single-shot SEBOLD with EPI readout, with all its known disadvantages, still is a very robust and time-efficient way to acquire SE-BOLD data.

An interesting new approach for SE-BOLD is using an echo planar time-resolved imaging (EPTI) readout (Wang et al., 2021). This technique promises the acquisition of SE-BOLD data, from which *T*_2_-weighted images without 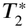 contamination can be reconstructed. While including these different variants is out of scope of the current study, these alternatives should be considered for future cortical-depth dependent studies that rely on *T*_2_-weighed imaging methods.

The determination of the PSF relies on the linearity and shift-invariance of the imaging system, which allows the description of the PSF as a convolution operation. However, this approach does not account for any spatial variations. Confounding factors, such as eye movements, may introduce spatially dependent variability. In addition, the cortical macrovasculature is sparse and heterogeneously distributed (Duvernoy, Delon, and Vannson, 1981), and the current study shows (see ***Figure 11***) that the local vascular geometry influences the spatial specificity of the BOLD signal. The estimated PSF therefore represents an effective PSF averaged over locations with very different macrovascular contributions within V1 rather than a local property of each cortical location. Thus, the current model is a simplification of the underlying mechanism.

Furthermore, our model assumes a simple Gaussian white noise term added to complex-valued time series, which allows us to describe the expected value of BOLD signal changes at stimulus frequency as a Rice distribution (see Appendix: Derivation of the modulation transfer function model). This approach is only strictly valid for the thermal noise limit (Triantafyllou et al., 2005) and, therefore, disregards physiological noise features and complexities due to parallel imaging and other multi-coil effects (Todd et al., 2017; Vizioli et al., 2021). In agreement with this, ***Figure 11*** showed that the noise level also depends on the distance to venous compartments, which points toward some remaining physiological noise contributions in the data.

Another confounding factor found in all studies that exploit the retinotopic organization of the visual cortex is the finite width and scatter of receptive fields (Hubel and Wiesel, 1974; Chaimow et al., 2018), i.e., neuronal activity already results in a blurred representation of the visual stimulus. The spatial specificity determined on the basis of receptive field properties cannot be disentangled from the estimated BOLD PSF. Therefore, width and scatter of receptive fields in V1 define an upper bound for the estimation of BOLD PSF. However, it is noted that this only affects the absolute estimates and not the comparison between GE- and SE-BOLD. A recent study analyzed the cortical depth dependence of pRFs (Fracasso, Petridou, and Dumoulin, 2016) in humans that showed a U-shaped profile across cortical depth, which is also expected based on neurophysiological findings (Chapin, 1986). The steady increase in PSF width found in the current study thus shows that the cortical profile is not just mirroring receptive field properties across cortical depth.

Furthermore, our estimates are specific to the employed stimulation paradigm. We used a phase-encoded design, which is a standard procedure for retinotopic mapping, with long runs and a relatively slow stimulation period (48 s) to induce well-defined traveling waves for an accurate estimation of the MTF. This results in a continuous stimulation of the visual cortex. As pointed out by Parkes et al., 2005, a large vein draining much of V1 can receive blood from an activated region at essentially every point in the rotation of the stimulus. Thus, its BOLD signal can remain elevated throughout the stimulation cycle. Consequently, our results may not directly translate to block designs or short event-related designs. Investigating this dependency is, however, beyond the scope of the current study.

In ***Supplementary Figure 6***, the large variability of parameter estimates between participants can be seen. During measurements, participants needed to fix their gaze on the central point for the whole time of data acquisition. No explicit task was given and we did not control for eye movements during measurements. Furthermore, GE- and SE-BOLD runs were acquired in separate sessions. Together, these factors limit our ability to control for behavioral differences in subject compliance across sessions, which might have contributed to the observed variability in signal amplitudes between sequences, sessions, and runs. Future studies could interleave the acquisition of both sequences within the same session to better control this confound.

Please note that the primary motivation for using a hierarchical Bayesian framework for data analysis was to obtain robust parameter estimates at the group level while acknowledging differences between subjects that cannot be explained by our MTF model.

In conclusion, the current study estimated and compared for the first time the BOLD PSF of GE- and SE-BOLD at 7 T at different cortical depths. The cortical profiles show that both acquisition techniques are similarly affected by macrovascular signal contributions that limit their spatial specificity.

## Appendix: Derivation of the modulation transfer function model

The modulation transfer function of the BOLD response (*MT F*_*BOLD*_) is estimated from traveling waves of neuronal activity in retinotopically organized cortex that were induced by rotating wedge stimuli. This section provides the derivation of the MTF model that was used to estimate the spatial specificity of the BOLD signal. Let *w*(*x, t*) be a traveling wave on the cortical surface that can be parameterized by cortical space *x* and time *t*

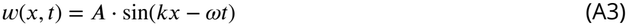

with spatial frequency *k* and temporal frequency *ω*. The temporal frequency only depends on the used stimulation frequency *ω*_*s*_, which was held constant for different number of wedges *ω* = *ω*_*s*_. The spatial frequency in units of cycles per mm depends on the applied stimulus (number of wedges) and the traveling distance on cortex (length of iso-eccentricity line) *k*_*ecc*_ = *N*_*wedge*_∕_2*D*_, where *N*_*wedge*_ is the number of wedges and *D* the traveling distance on one hemisphere. The factor 2 is included to get the full distance for one stimulus rotation (Parkes et al., 2005). Without loss of generality, we set *A* = 1. The traveling wave on the cortical surface along one iso-eccentricity line has then the following form

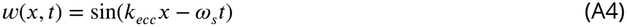

Neuronal activity *n*(*x, t*) is assumed to be positive in the range [0, 1] with no and maximal activity indicated by 0 and 1, respectively.

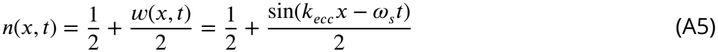

We assume that the fMRI signal can be modeled as a shift-invariant linear process, which allows to describe the measured response as convolution with a spatiotemporal point spread function kernel *PSF*_*BOLD*_(*x, t*) (Parkes et al., 2005). The measured BOLD activity on the cortex *b*(*x, t*) is then given by

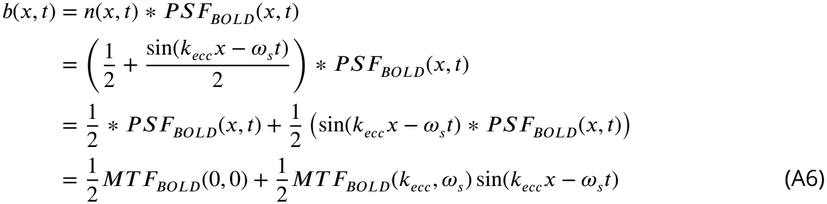

In equation A6, we used the Convolution theorem to express the convolution with the *PSF*_*BOLD*_(*x, t*) kernel by point-wise multiplication with its Fourier transform, the modulation transfer function *MT F*_*BOLD*_(*k, ω*). In an actual fMRI experiment, we acquire a complex-valued time series with *N* measurements separated by the repetition time TR. The time series of one voxel at position *x*_0_ is then given by

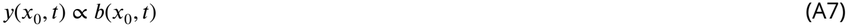

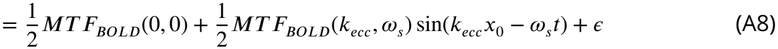

We assume stimulus-independent Gaussian white noise *ϵ ∼ N*(0, *σ*^2^) that is added to the time series. The noise level will lead to a non-zero baseline level in magnitude images that are used for analysis (Vizioli et al., 2021). Therefore, it has to be incorporated in the model to get an unbiased estimation of the *MT F*_*BOLD*_, especially when comparing acquisition techniques like GE- and SEBOLD that are affected by different signal-to-noise (SNR) penalties. We perform a discrete Fourier analysis in time and extract the absolute value of the Fourier component *c*_*s*_(*k*_*ecc*_) at stimulation frequency *ω*_*s*_.

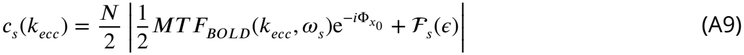

The first term results in an impulse response at stimulation frequency whose magnitude is scaled by the number of time points ^*N*^∕_2_ and phase is shifted by 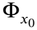 . The second term is the Fourier transform of the noise term. We now focus on the expected value of the Fourier component.

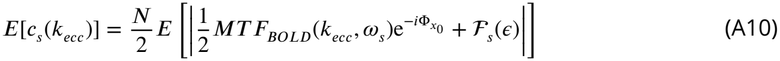

The underlying distribution of the expected value at *k*_*ecc*_ follows a Rice distribution (Gudbjartsson and Patz, 1995). Using the expression for the first moment of the Rice distribution (Park Jr., 1961), the expected value of the absolute value of the Fourier component at simulation frequency is given by

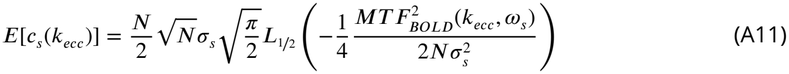

with the fractional Laguerre polynomial *L*_1∕2_ (Abramowitz and Stegun, 1964). For *MT F*_*BOLD*_(*k*_*ecc*_, *ω*_*s*_), we assume a Gaussian distribution.

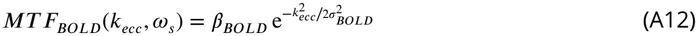

The final spread of the BOLD response is evaluated by the full width at half maximum (FWHM) of the associated *PSF*_*BOLD*_.

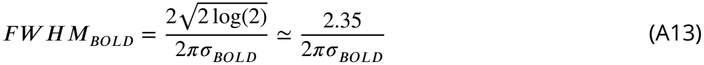

The final *MT F*_*BOLD*_ model has the three free parameters *β*_*BOLD*_ (amplitude), *F W HM*_*BOLD*_ (width) and *σ*_*s*_ (baseline offset).

## Acknowledgments

The research leading to these results has received funding from the European Research Council under the European Union’s Seventh Framework Program (FP7/2007-2013) / ERC grant agreement n° 616905 and from the Deutsche Forschungsgemeinschaft (DFG, German Research Foundation) — project n° 347592254 (WE 5046/4-2 and/or KI 1337/2-2). Nikolaus Weiskopf has received funding from the European Union’s Horizon 2020 research and innovation program under the grant agreement n° 681094 and from the BMBF (01EW1711A & B) in the framework of ERA-NET NEURON. We thank the University of Minnesota Center for Magnetic Resonance Research for the provision of the multiband EPI sequence software. We thank Johanna Bergmann for providing the scripts for population receptive field modeling.

## Author contributions

Daniel Haenelt, Conceptualization, Methodology, Software, Formal analysis, Investigation, Data curation, Writing - original draft preparation, Writing - review & editing, Visualization; Robert Trampel, Investigation, Writing - review & editing, Supervision; Denis Chaimow, Conceptualization, Methodology, Writing - review & editing; Amir Shmuel, Methodology, Writing - review & editing; Martin I. Sereno, Methodology, Software, Writing - review & editing; Nikolaus Weiskopf, Conceptualization, Resources, Writing - review & editing, Supervision, Project administration, Funding acquisition

## Supplementary Materials

**Supplementary Figure 1.**
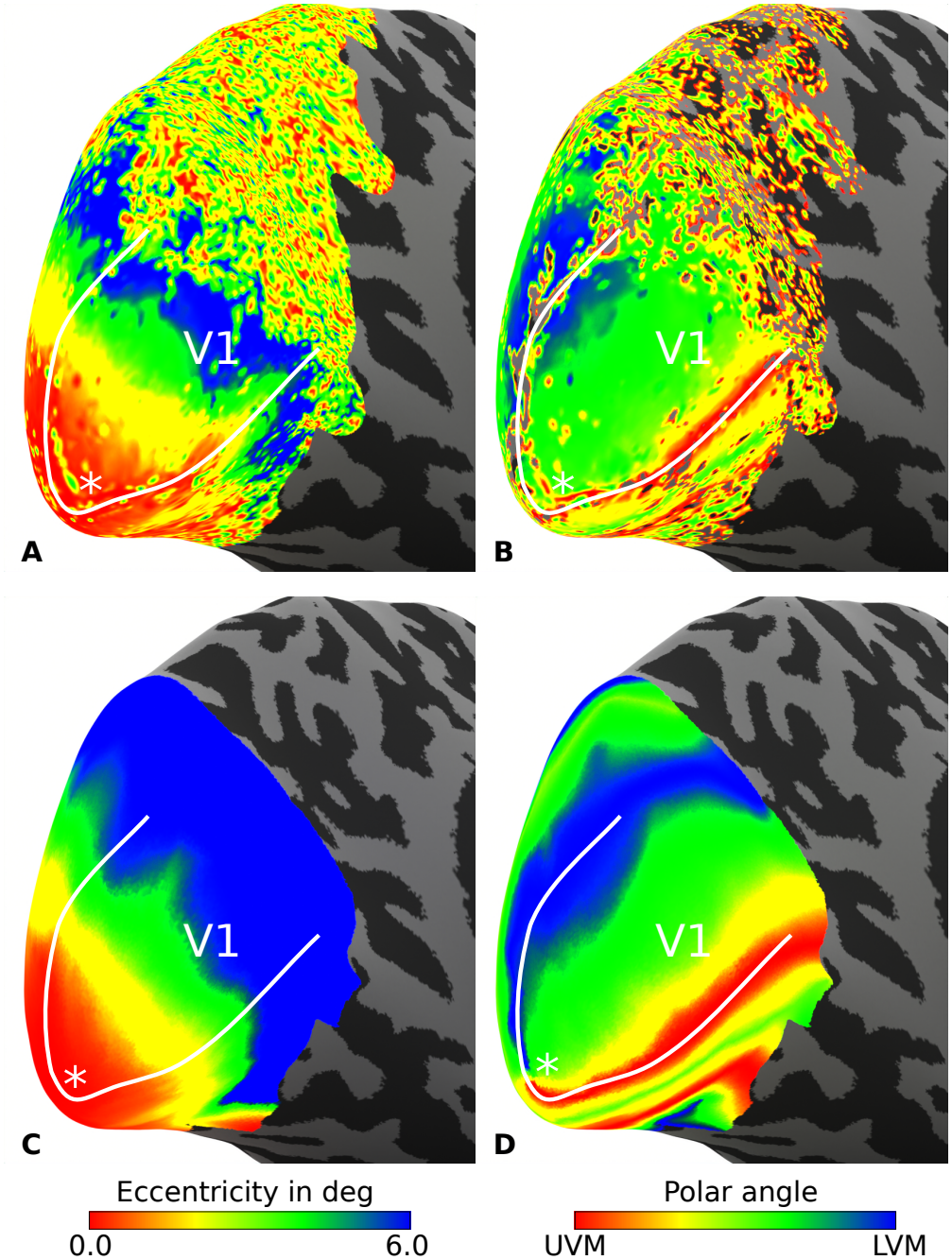
Typical retinotopic maps. For one representative participant (subject 6), maps for eccentricity and polar angle are shown in **A** and **B**, respectively. These maps are based on voxel-wise analysis of fMRI data, which are prone to measurement noise that hinders an accurate assessment of cortical distances (see ***Figure 6***). Therefore, we used the method by Benson and Winawer, 2018 that takes measured retinotopy data as prior information to compute a quasi-conformal mapping from a retinotopy template to the subject anatomy. Resulting eccentricity and polar angle maps are shown in **C** and **D**, respectively. Analysis was restricted to data sampled at mid-cortical depth. Solid white lines indicate the V1/V2 border that was based on polar angle maps. White asterisks mark the location of the foveal representation.

**Supplementary Figure 2.**
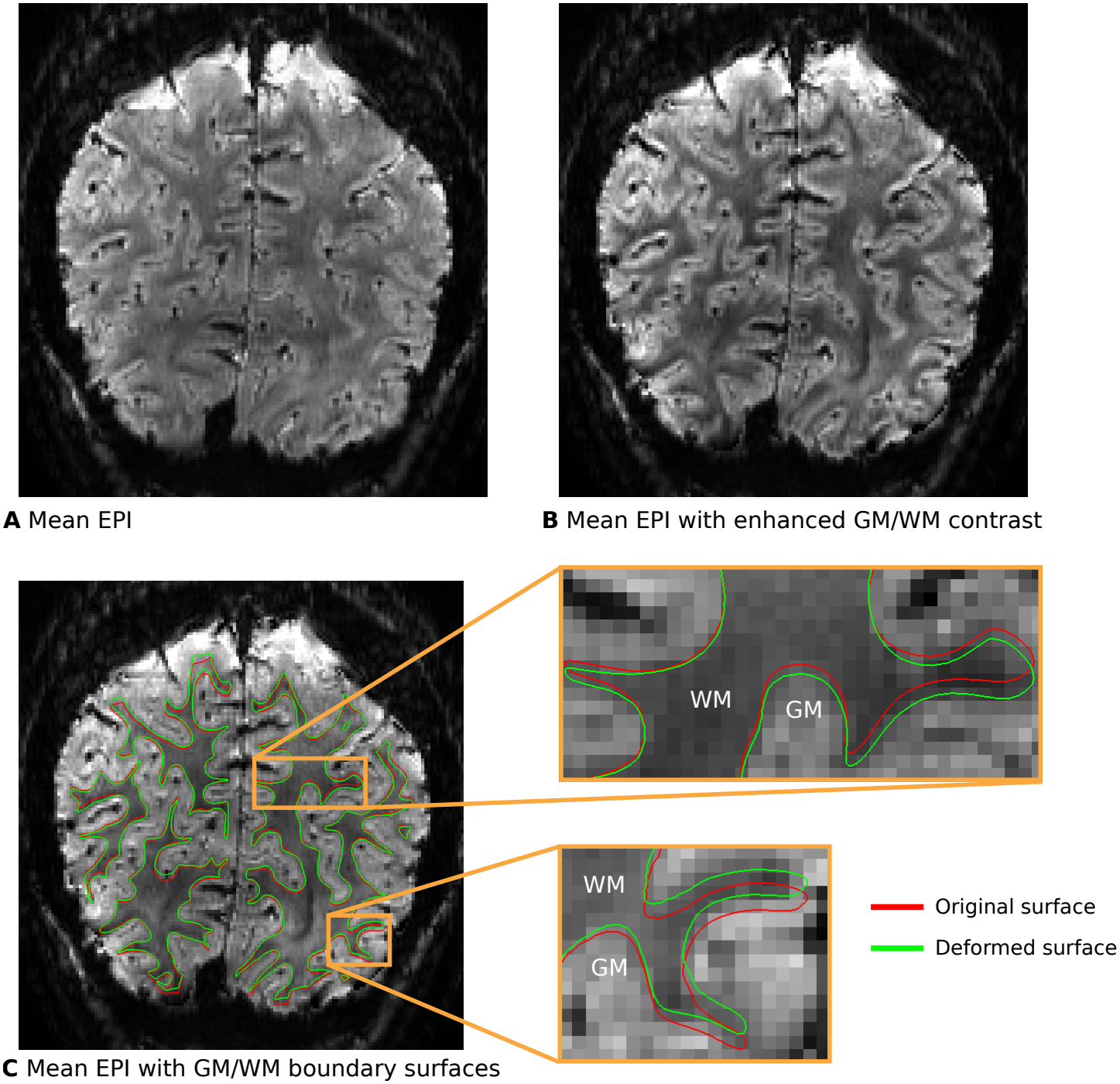
Exemplary illustration of the gradient-boundary based (GBB) method. To improve the alignment of cortical boundary surfaces with the GM/WM border found in functional images, we first transformed surfaces to the space of functional time series (see Data preprocessing for further details). Then, the surfaces were moved along the image gradient calculated from the temporal mean of the functional time series until they hit the GM/WM border of the distorted functional image. This method is implemented in the GBB package (0.1.6, https://pypi.org/project/gbb/). In **A**, the temporal mean of one functional time series from subject 1 is shown in coronal view. **B** To enhance the contrast of the GM/WM border, we followed the method suggested in Fracasso, Petridou, and Dumoulin, 2016 and weighted the magnitude image by its phase (both provided by the online reconstruction on the scanner), as typically done in susceptibility-weighted imaging methods. More precisely, the magnitude time series was first realigned using AFNI (19.1.05, https://afni.nimh.nih.gov/) (Cox, 1996). Each volume of the phase time series was then individually unwrapped utilizing the method of Abdul-Rahman et al., 2005 implemented in Nighres (1.2.0, https://pypi.org/project/nighres/) (Huntenburg, Steele, and Bazin, 2018). Computed motion parameters were then applied to the unwrapped phase time series. The temporal mean of magnitude and phase data were then derived, and the phase data was further thresholded and normalized. Finally, the contrast of the magnitude data was enhanced by weighting each voxel by its contrast-reversed phase data. **C** shows surfaces before (red) and after (green) alignment with the GBB method. Surfaces coincide better with the GM/WM border found in functional images after alignment.

**Supplementary Figure 3.**
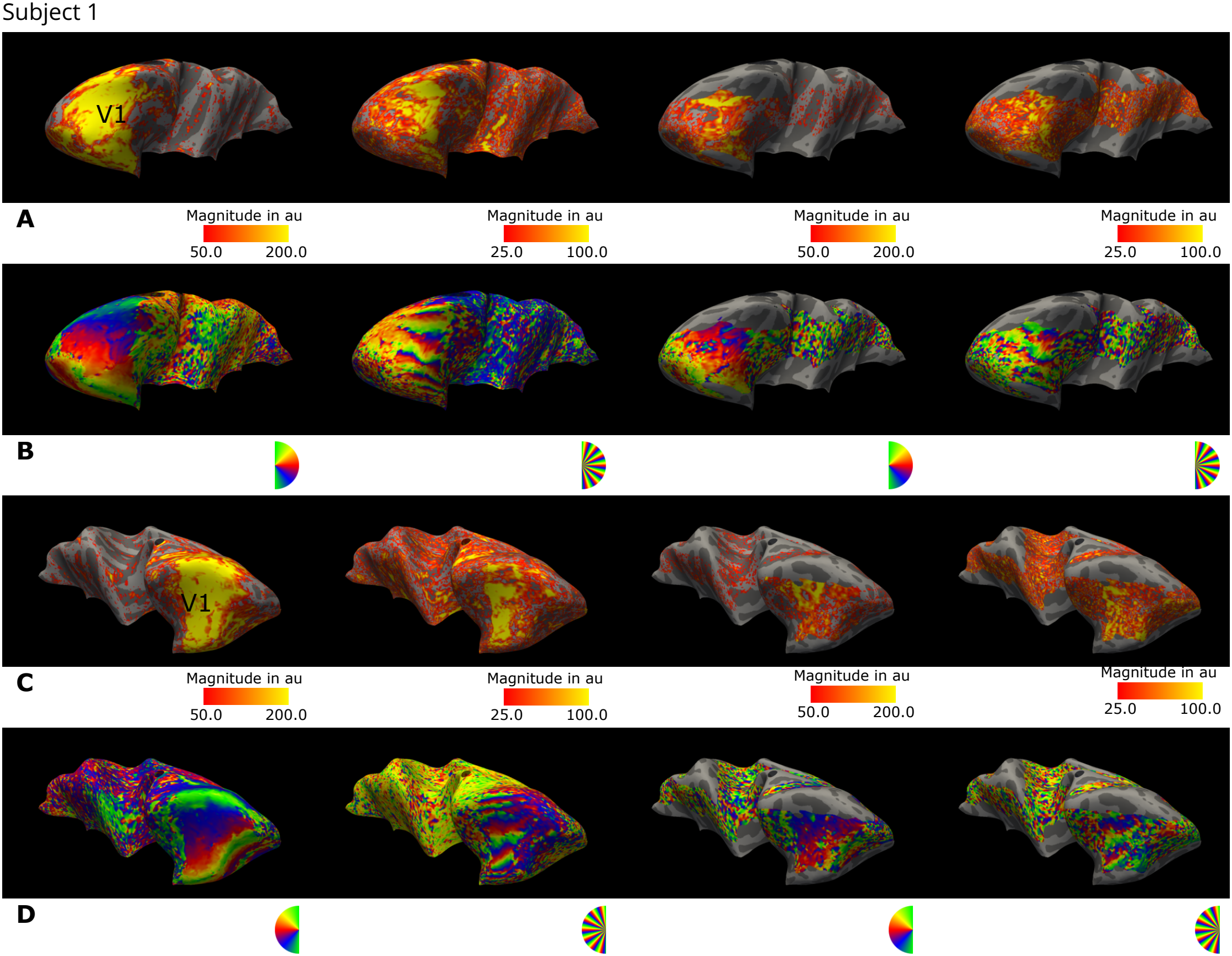
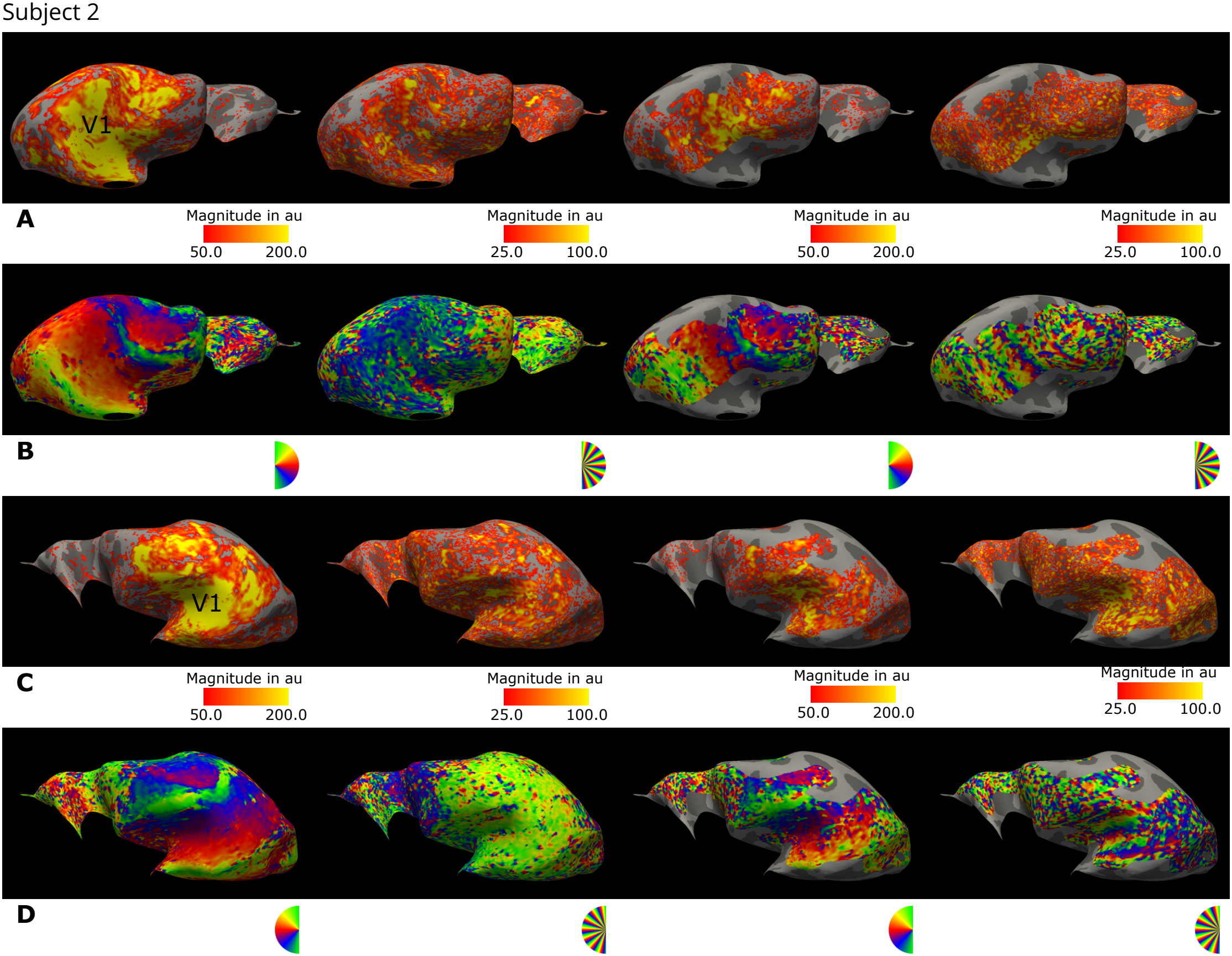
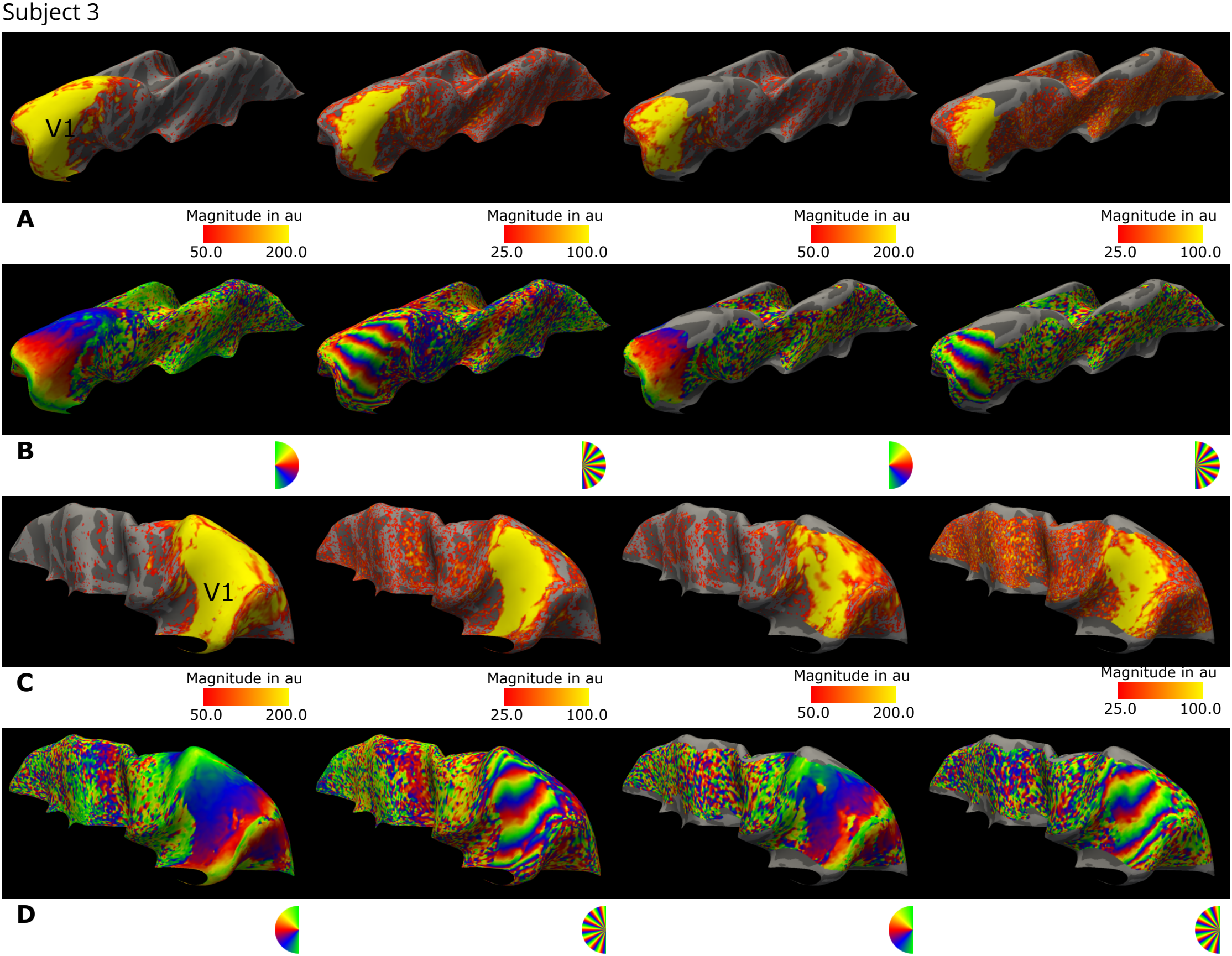
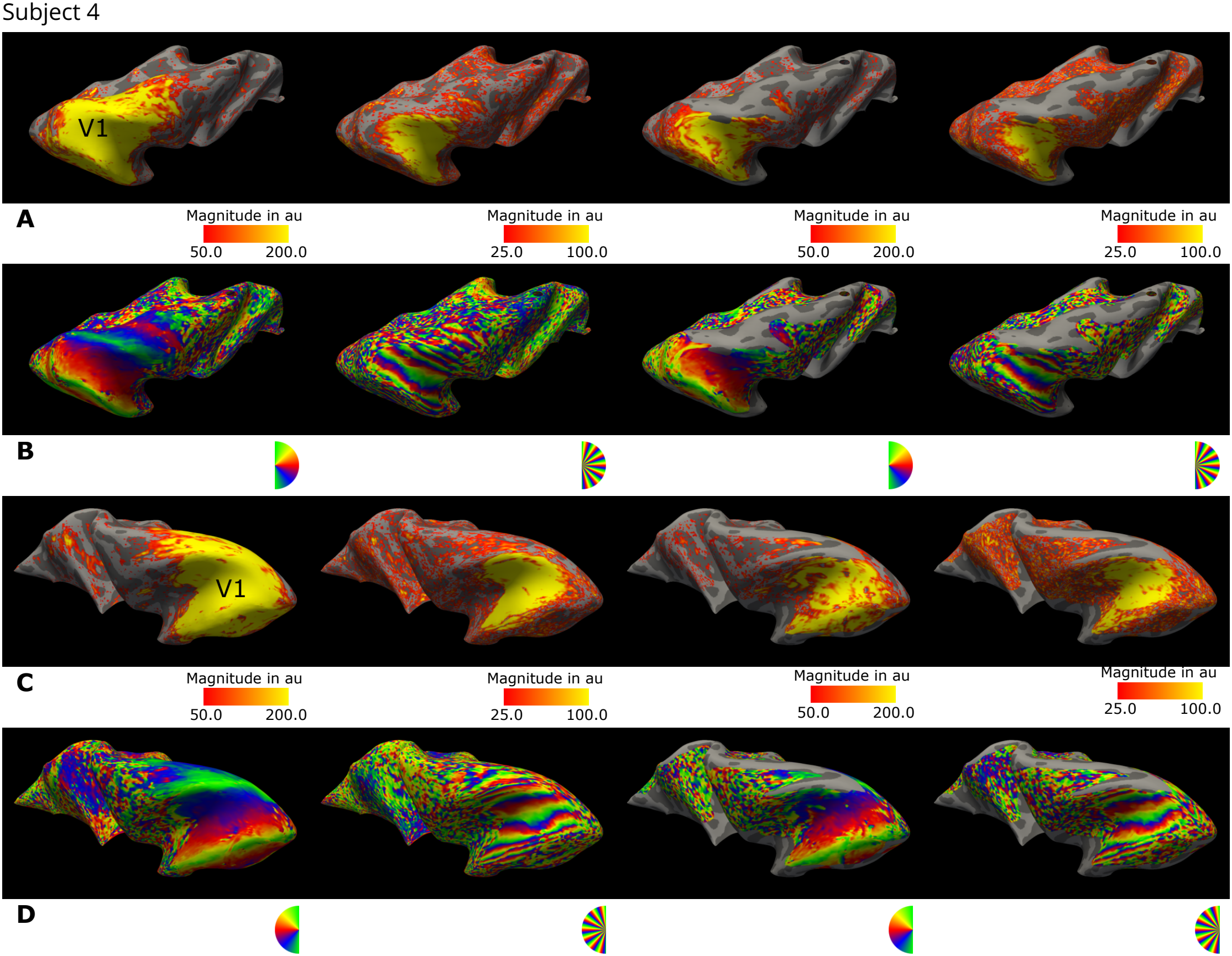
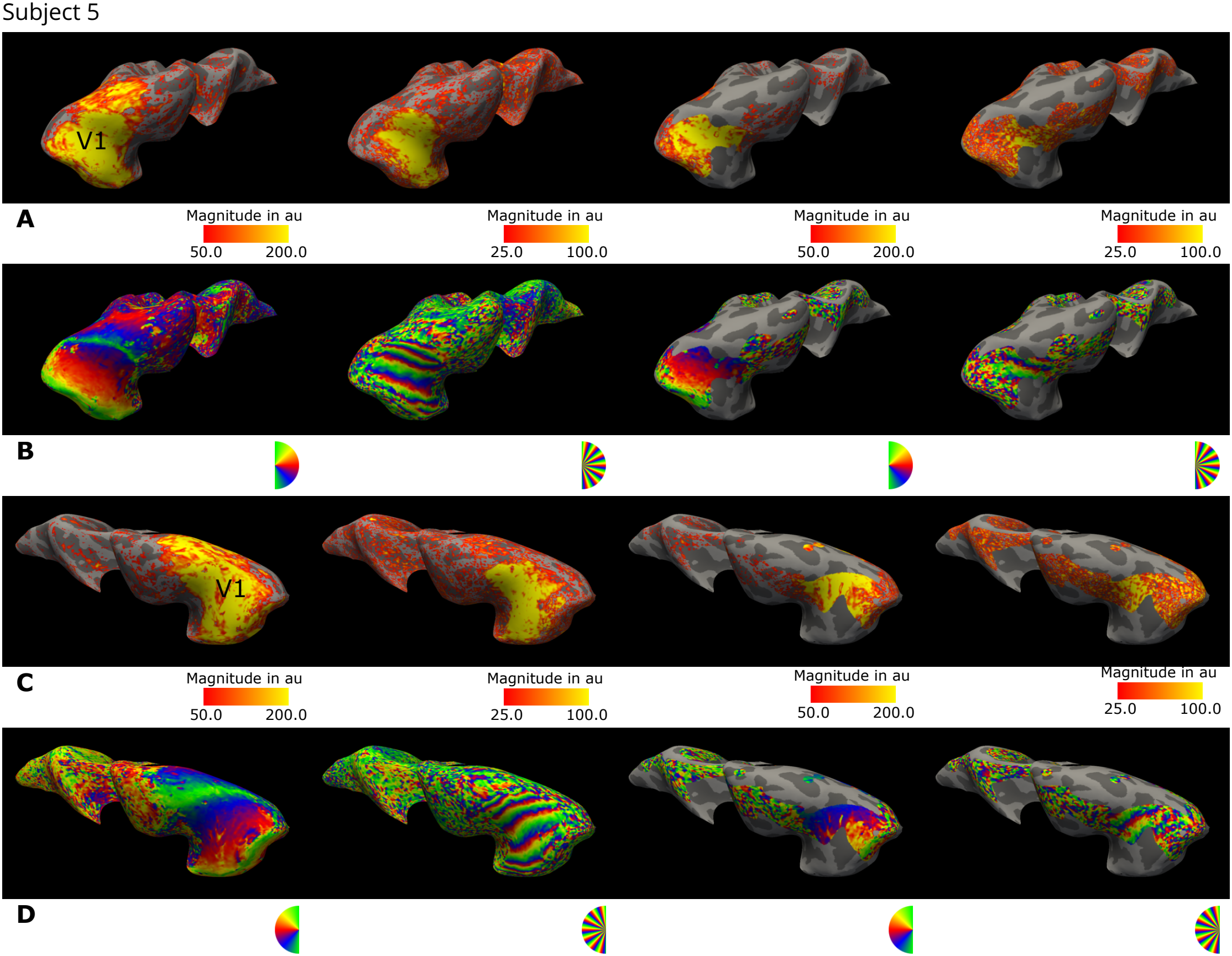
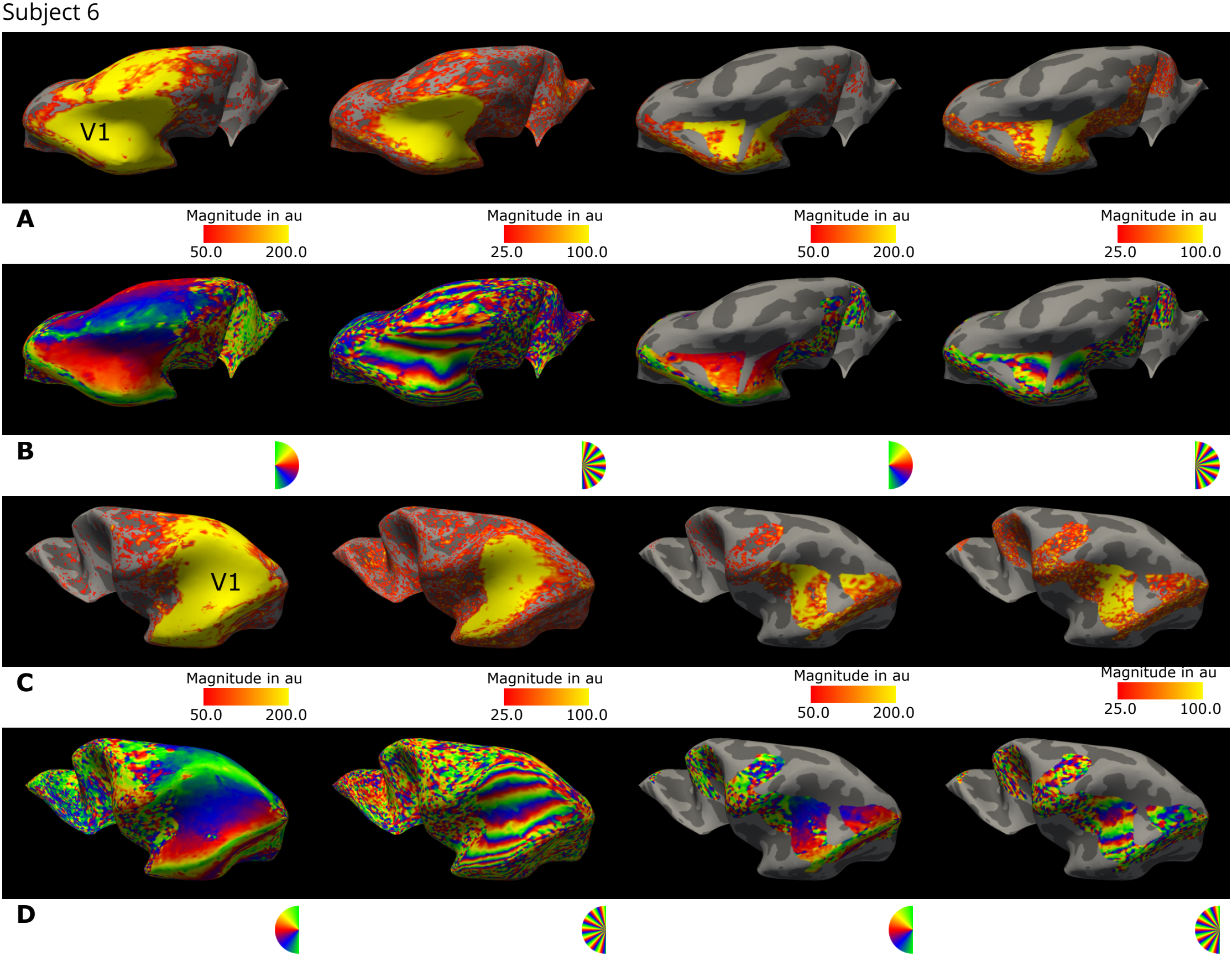
Phase and magnitude responses at stimulus frequency. For a single participant, the magnitude **(A, B)** and phase **(C, D)** responses are shown for the left and right hemispheres, respectively. Data were sampled on the mid-cortical surface, averaged across two sessions, and only show regions covered by the functional acquisitions. Projected data represent the average of two runs from separate sessions. The occipital pole in this representation is located on the left and right for the left and right hemispheres, respectively. Columns 1–2 display GE-BOLD data, while columns 3–4 show SE-BOLD data. Note the smaller coverage in SE-BOLD acquisitions. For each sequence, data from runs with 2 and 14 wedges are shown in columns (1, 3) and (2, 4), respectively.

**Supplementary Figure 4.**
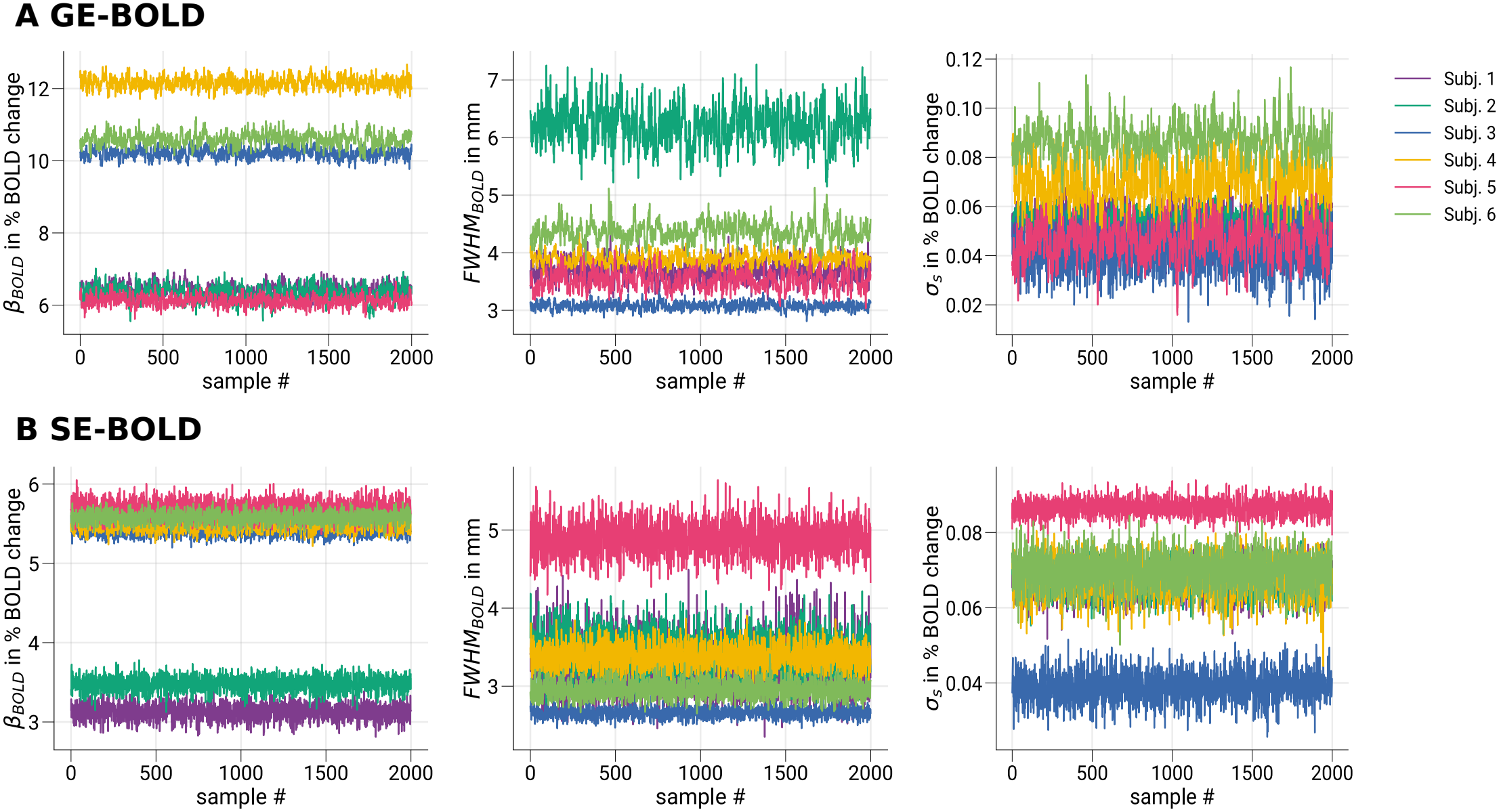
Exemplary single-subject trace plots for convergence diagnostics of Markov chain Monte Carlo sampling. In **A** and **B**, exemplary single-subject trace plots for sampled parameter estimates are shown for GE- and SE-BOLD, respectively. No significant trends are visually noticeable within traces of single participants, indicating reached convergence (stationarity). In contrast, larger differences can be seen between participants.

**Supplementary Figure 5.**
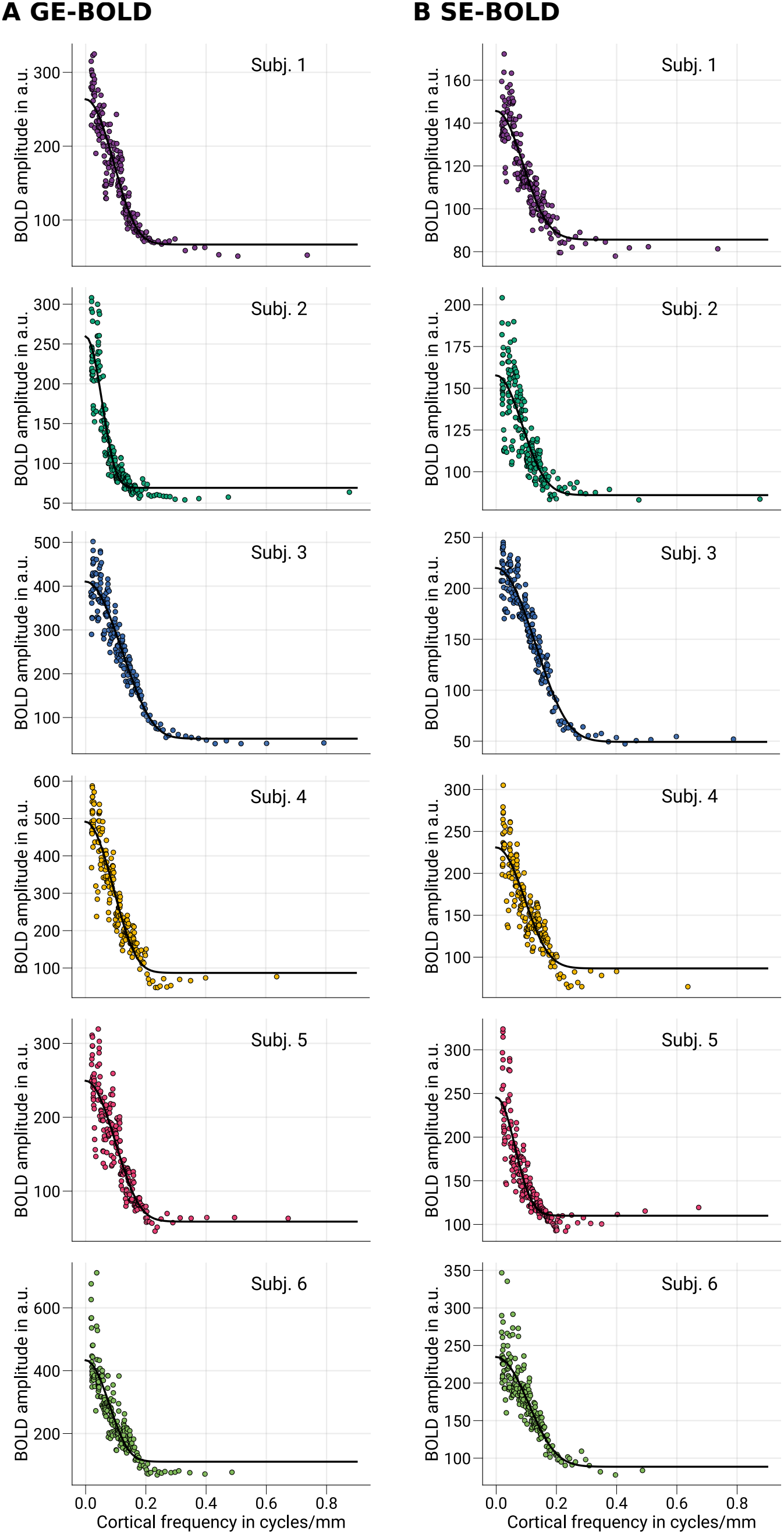
Exemplary single-subject MTF curves for GE- and SE-BOLD. In **A** and **B**, single-subject MTF model fits are shown for GE- and SE-BOLD measurements from data sampled at mid-cortical depth. Data points show BOLD amplitudes binned by spatial frequency (*n* = 200). Black lines indicate MTF curves based on the median of parameter estimates at the subject level.

**Supplementary Figure 6.**
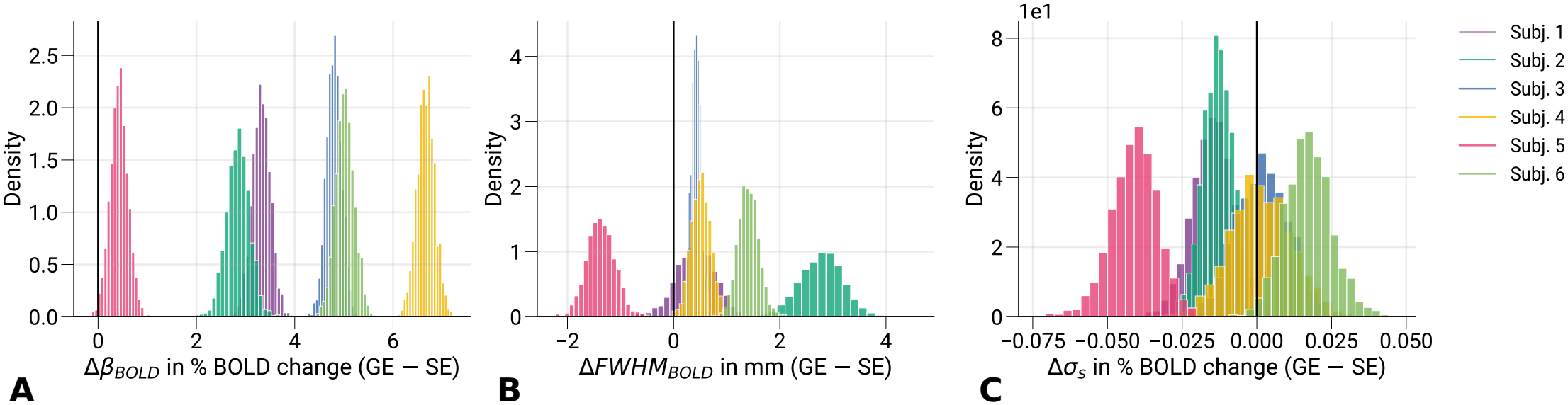
Differences between GE- and SE-BOLD parameter estimates. Differences between GE- and SE-BOLD parameter estimates are shown for **A** Δ*β*_*BOLD*_, **B** Δ*F W HM*_*BOLD*_ and **C** Δ*σ*_*s*_ at the subject level from data sampled at mid-cortical depth. Vertical black lines indicate the difference of zero. As expected, all participants show larger amplitudes (**A**) for GE-compared to SE-BOLD. The FWHM (**B**) similarly is broader for GE-BOLD for all but one participant (subject 5). Interestingly, this participant also had the smallest amplitude differences in **A**. Since the GE- and SE-BOLD data were acquired in separate sessions on different days, this discrepancy can have other reasons (e.g., differences in attentional states). Note that we used Bayesian hierarchical modeling to account for these residual differences at the population level. The baseline offset (**C**) shows no conclusive differences at the subject level. Comparisons of GE- and SE-BOLD parameter estimates at the population level are shown in ***Figure 8***.

