## Supplemental Data 1 for "Spatial specificity of the functional gradient echo and spin echo BOLD signal across cortical depth at 7 T"

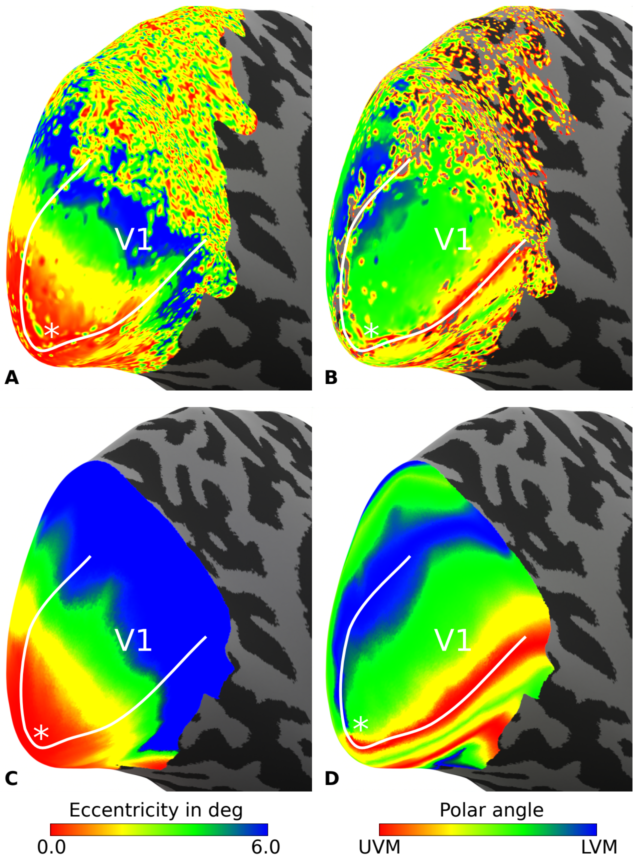


**Supplementary Figure 1. Typical retinotopic maps.** For one representative participant (subject 6), maps for eccentricity and polar angle are shown in **A** and **B**, respectively. These maps are based on voxel-wise analysis of fMRI data, which are prone to measurement noise that hinders an accurate assessment of cortical distances (see ***Figure 6***). Therefore, we used the method by Benson and Winawer, 2018 that takes measured retinotopy data as prior information to compute a quasi-conformal mapping from a retinotopy template to the subject anatomy. Resulting eccentricity and polar angle maps are shown in **C** and **D**, respectively. Analysis was restricted to data sampled at mid-cortical depth. Solid white lines indicate the V1/V2 border that was based on polar angle maps. White asterisks mark the location of the foveal representation.

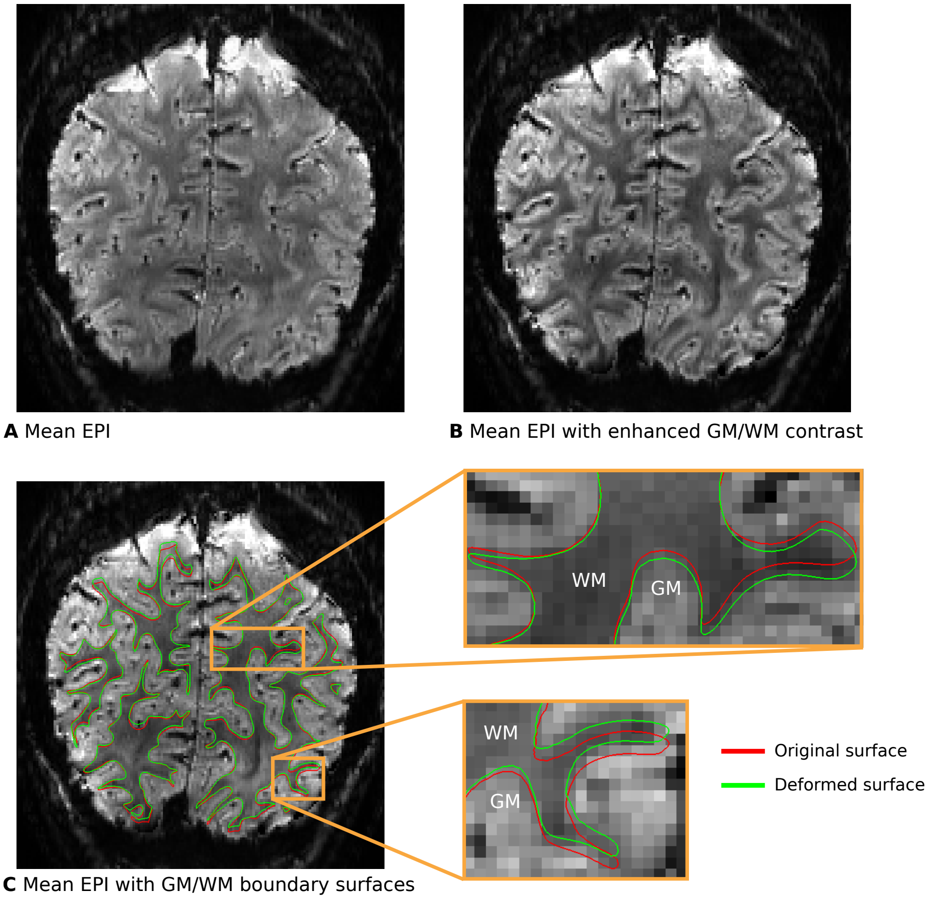


**Supplementary Figure 2. Exemplary illustration of the gradient-boundary based (GBB) method.** To improve the alignment of cortical boundary surfaces with the GM/WM border found in functional images, we first transformed surfaces to the space of functional time series (see Data preprocessing for further details). Then, the surfaces were moved along the image gradient calculated from the temporal mean of the functional time series until they hit the GM/WM border of the distorted functional image. This method is implemented in the GBB package (0.1.6, https://pypi.org/project/gbb/). In **A**, the temporal mean of one functional time series from subject 1 is shown in coronal view. **B** To enhance the contrast of the GM/WM border, we followed the method suggested in Fracasso, Petridou, and Dumoulin, 2016 and weighted the magnitude image by its phase (both provided by the online reconstruction on the scanner), as typically done in susceptibility-weighted imaging methods. More precisely, the magnitude time series was first realigned using AFNI (19.1.05, https://afni.nimh.nih.gov/) (Cox, 1996). Each volume of the phase time series was then individually unwrapped utilizing the method of Abdul-Rahman et al., 2005 implemented in Nighres (1.2.0, https://pypi.org/project/nighres/) (Huntenburg, Steele, and Bazin, 2018). Computed motion parameters were then applied to the unwrapped phase time series. The temporal mean of magnitude and phase data were then derived, and the phase data was further thresholded and normalized. Finally, the contrast of the magnitude data was enhanced by weighting each voxel by its contrast-reversed phase data. **C** shows surfaces before (red) and after (green) alignment with the GBB method. Surfaces coincide better with the GM/WM border found in functional images after alignment.

Subject 1

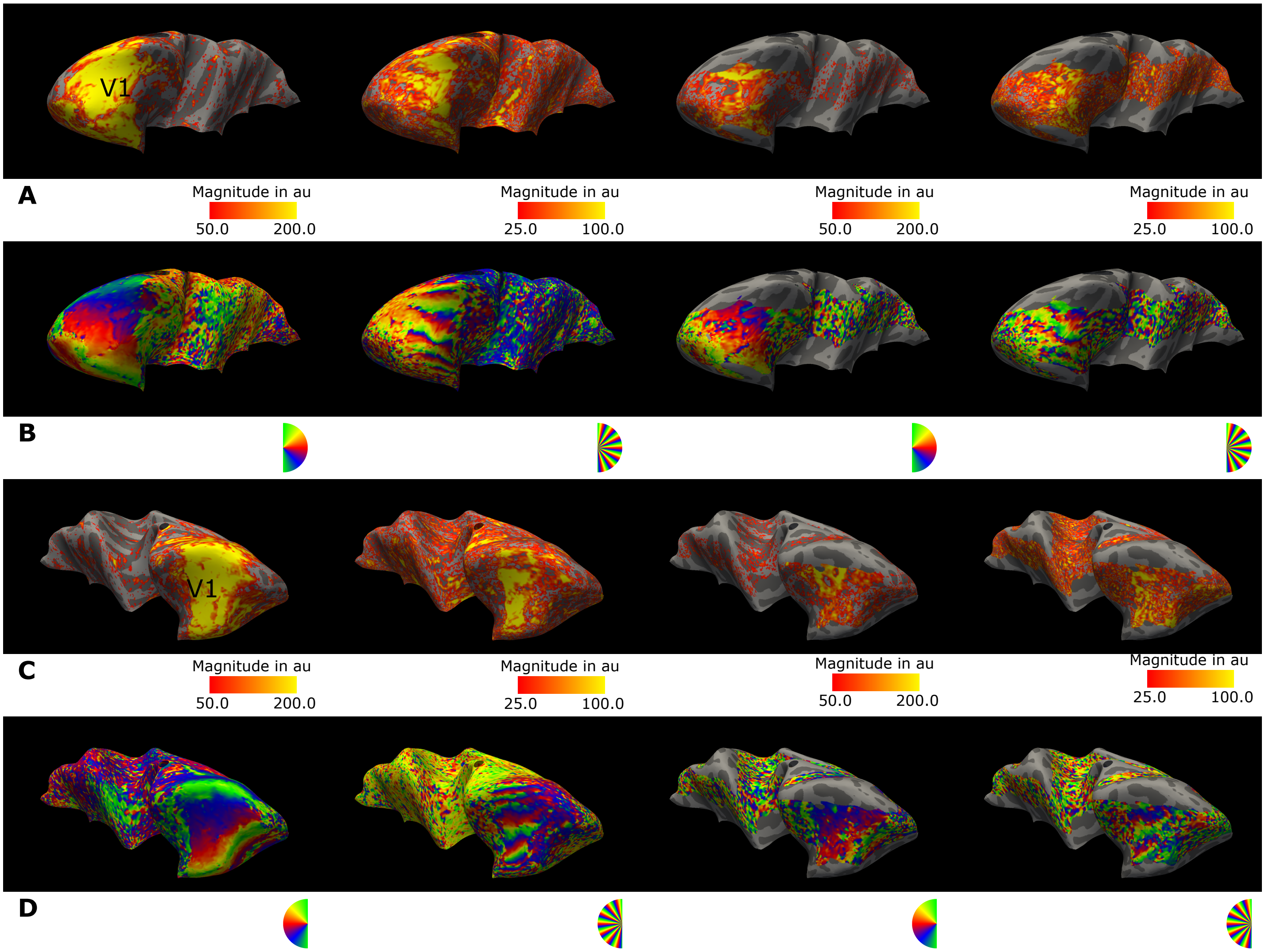


**Supplementary Figure 3. Phase and magnitude responses at stimulus frequency.** For a single participant, the magnitude **(A, B)** and phase **(C, D)** responses are shown for the left and right hemispheres, respectively. Data were sampled on the mid-cortical surface, averaged across two sessions, and only show regions covered by the functional acquisitions. Projected data represent the average of two runs from separate sessions. The occipital pole in this representation is located on the left and right for the left and right hemispheres, respectively. Columns 1–2 display GE-BOLD data, while columns 3–4 show SE-BOLD data. Note the smaller coverage in SE-BOLD acquisitions. For each sequence, data from runs with 2 and 14 wedges are shown in columns (1, 3) and (2, 4), respectively.

Subject 2

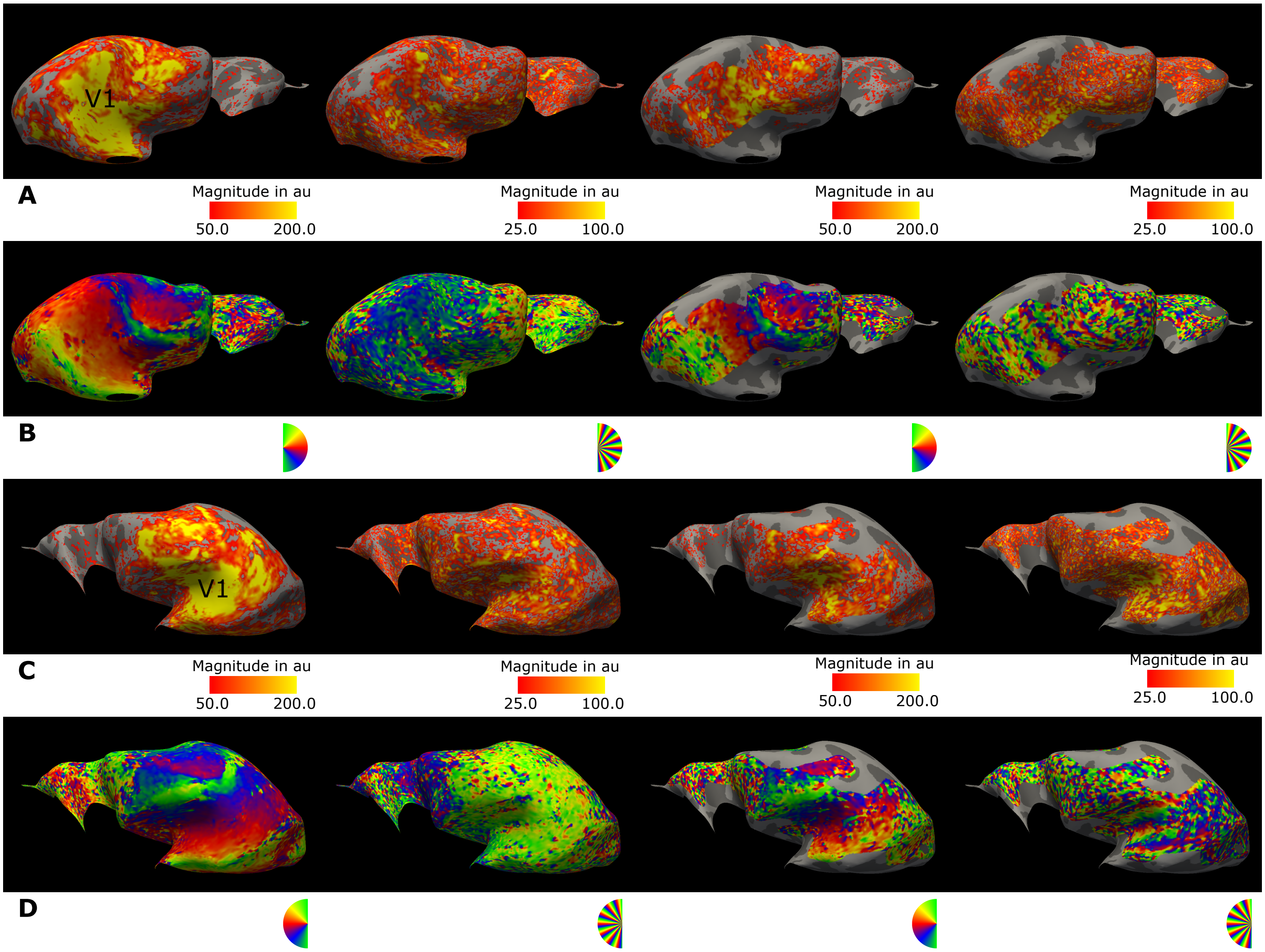


**Supplementary Figure 3.** (continued)

Subject 3

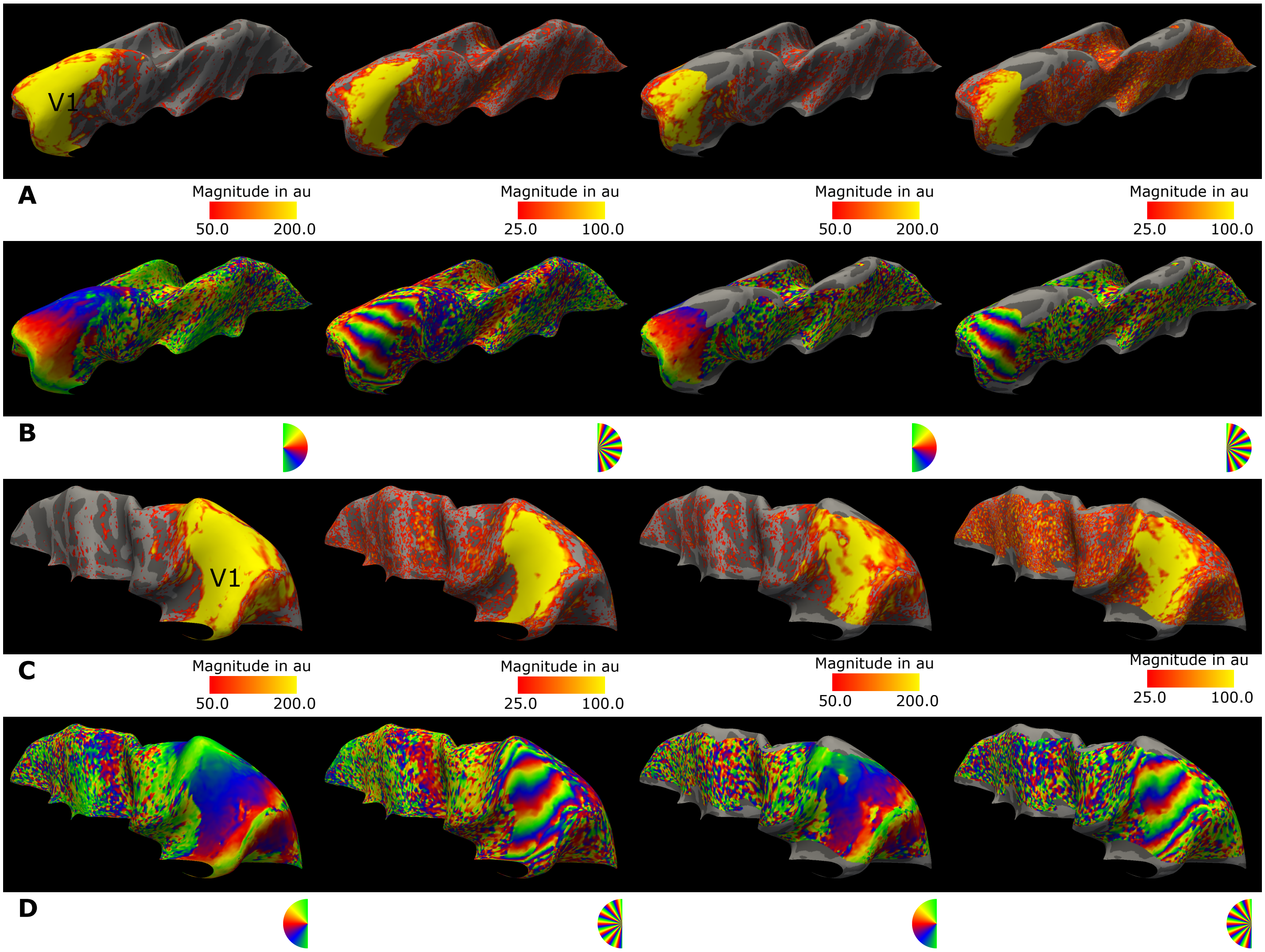


**Supplementary Figure 3.** (continued)

Subject 4

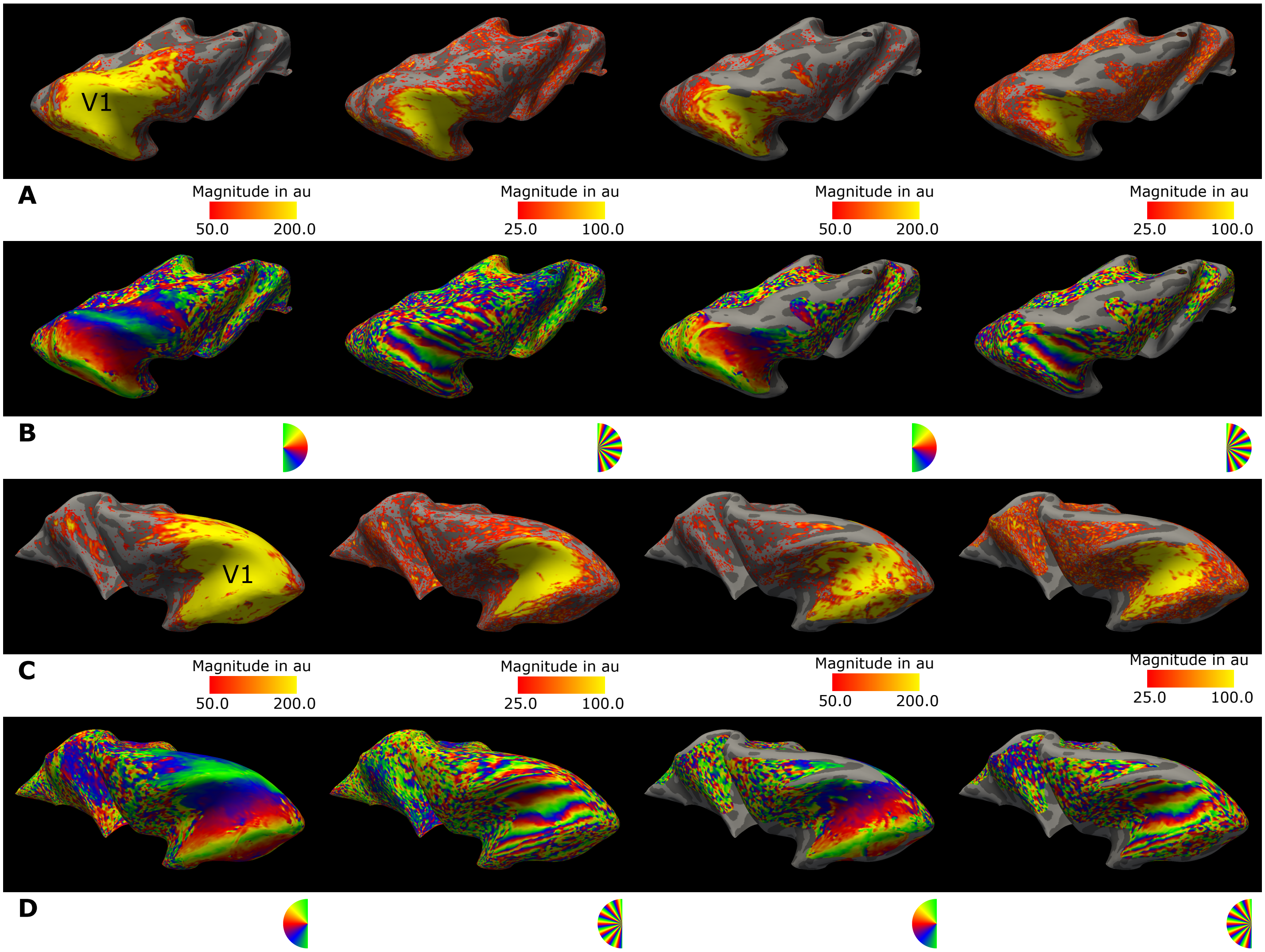


**Supplementary Figure 3.** (continued)

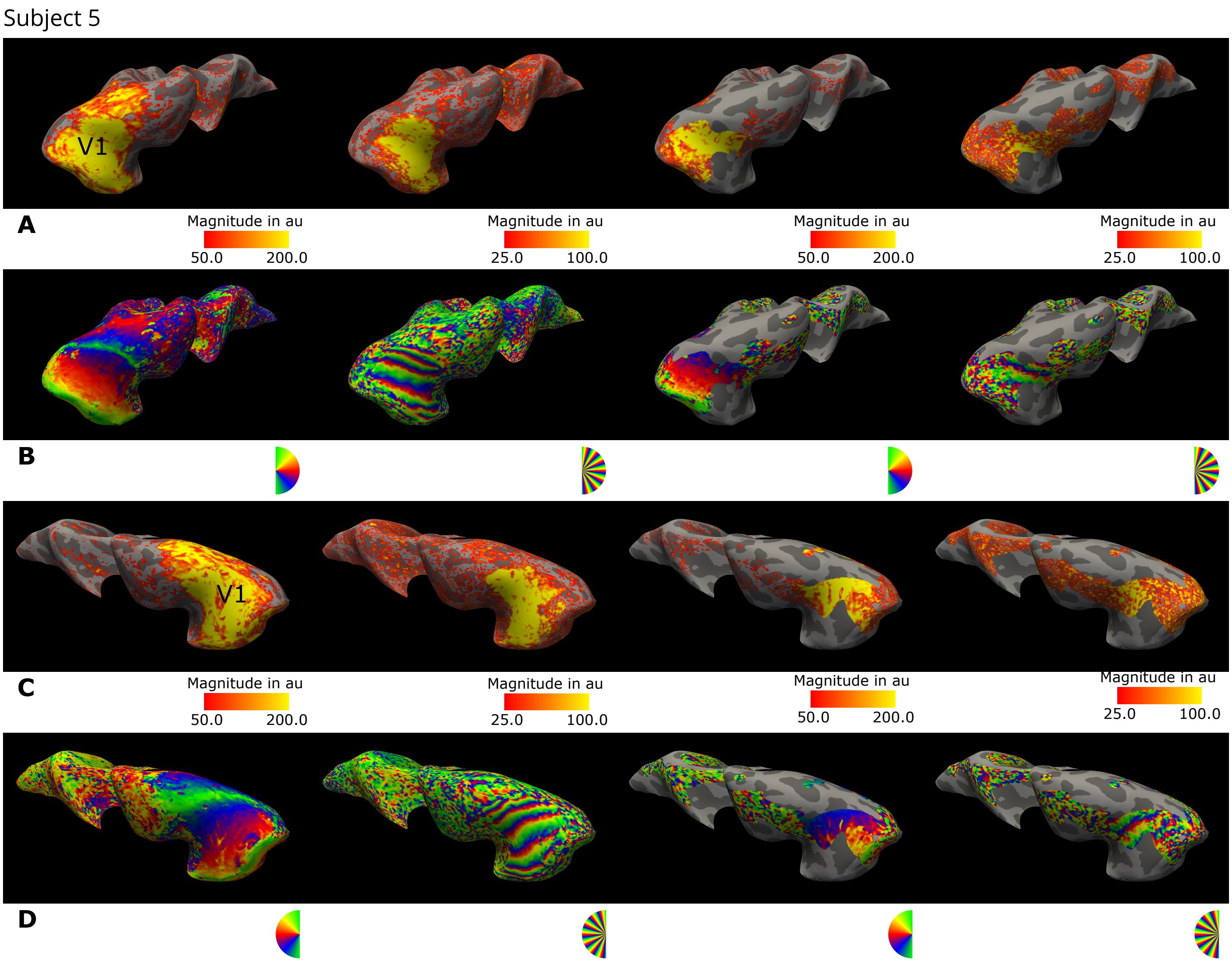


**Supplementary Figure 3.** (continued)

Subject 6

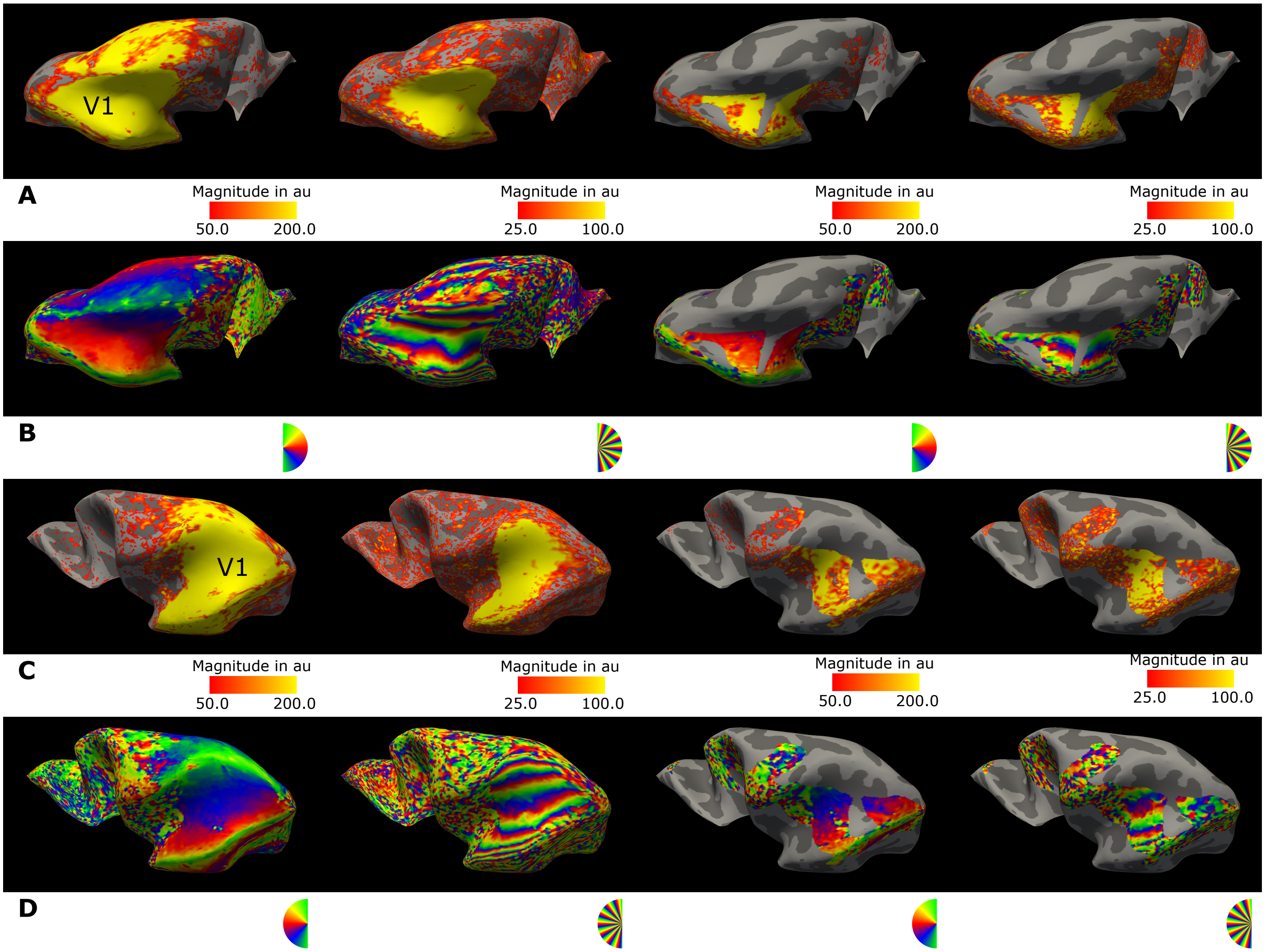


**Supplementary Figure 3.** (continued)

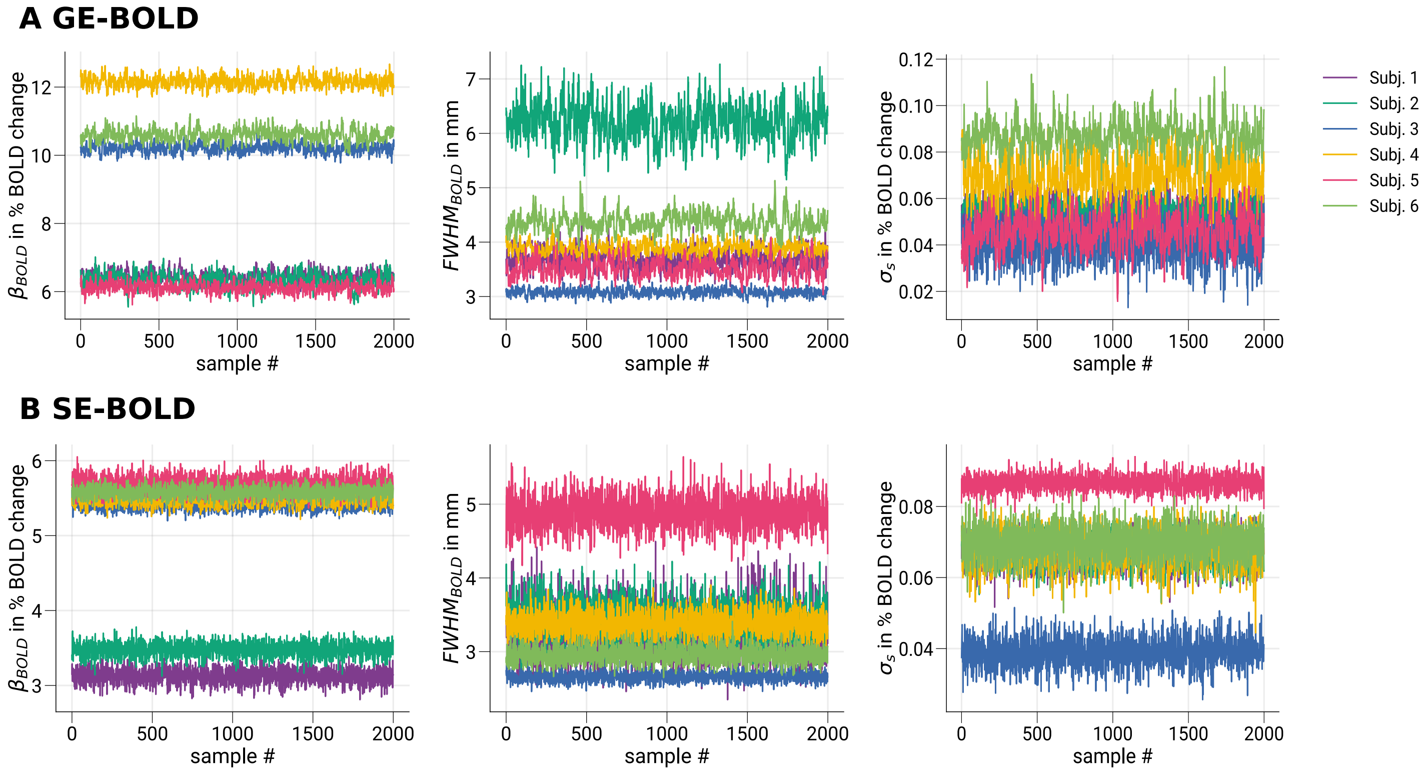


**Supplementary Figure 4. Exemplary single-subject trace plots for convergence diagnostics of Markov chain Monte Carlo sampling.** In **A** and **B**, exemplary single-subject trace plots for sampled parameter estimates are shown for GE- and SE-BOLD, respectively. No significant trends are visually noticeable within traces of single participants, indicating reached convergence (stationarity). In contrast, larger differences can be seen between participants.

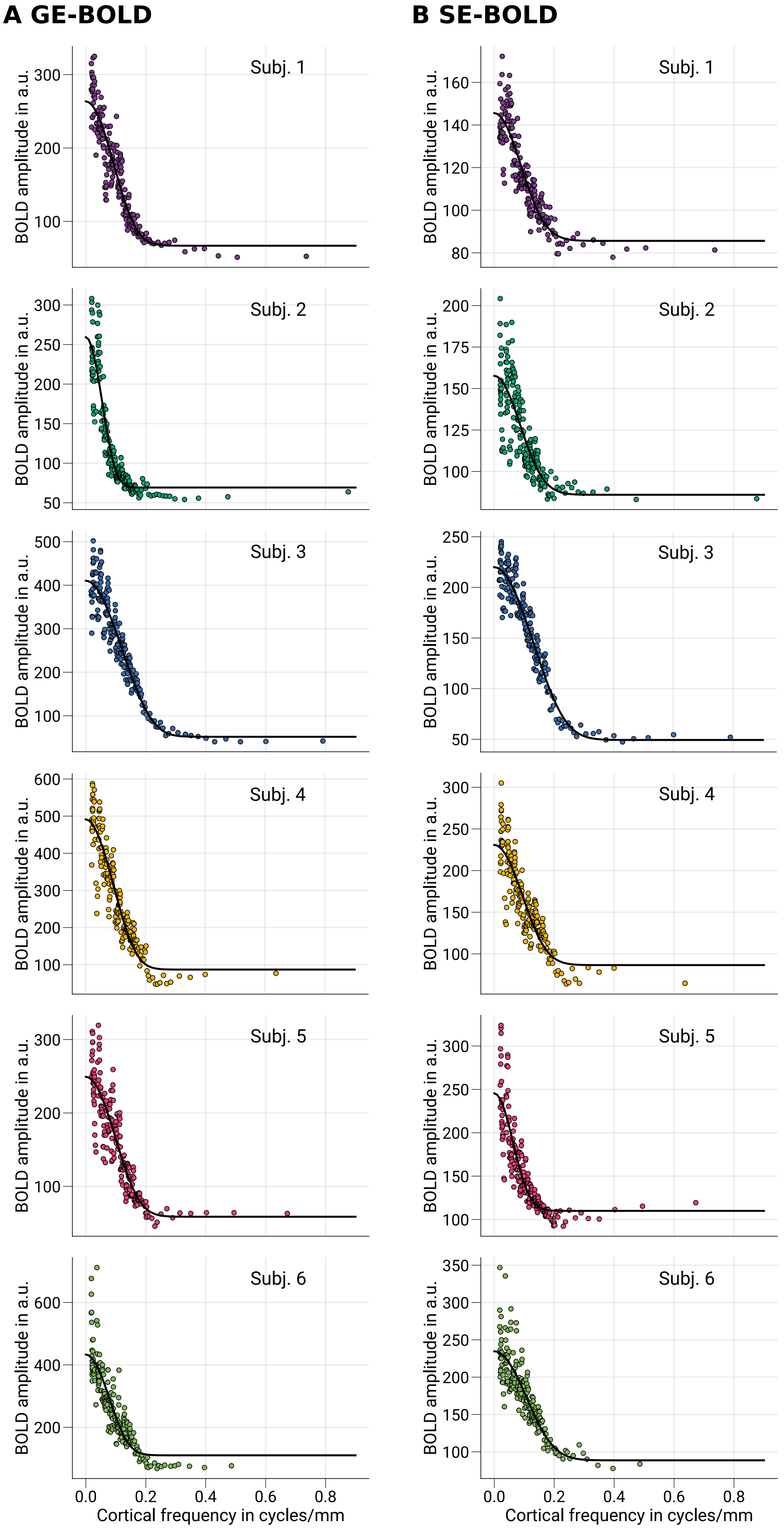


**Supplementary Figure 5.** (Caption next page.)

**Supplementary Figure 5. Exemplary single-subject MTF curves for GE- and SE-BOLD.** In **A** and **B**, single-subject MTF model fits are shown for GE- and SE-BOLD measurements from data sampled at mid-cortical depth. Data points show BOLD amplitudes binned by spatial frequency ($n = 200$). Black lines indicate MTF curves based on the median of parameter estimates at the subject level.

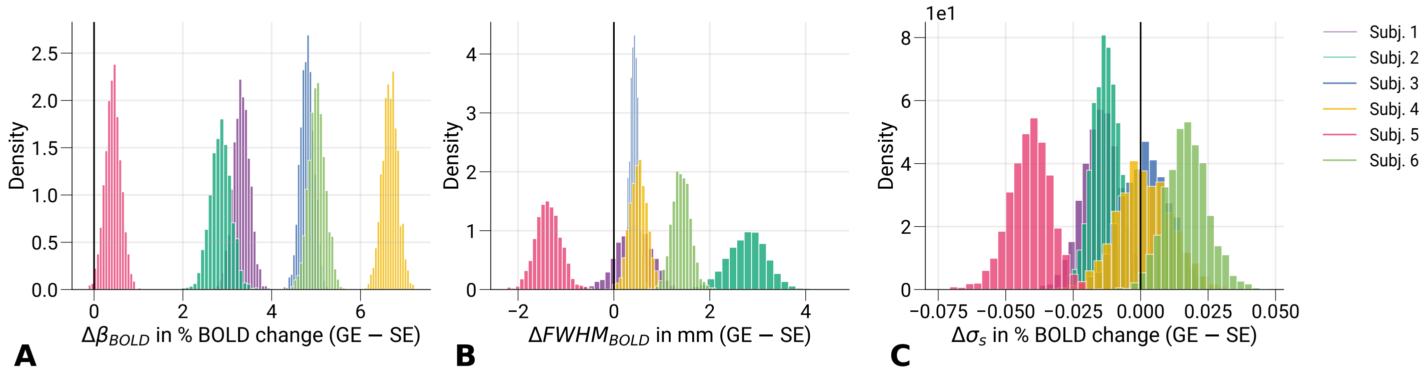


**Supplementary Figure 6. Differences between GE- and SE-BOLD parameter estimates.** Differences between GE- and SE-BOLD parameter estimates are shown for **A** $\Delta\beta_{BOLD}$, **B** $\Delta FWHM_{BOLD}$ and **C** $\Delta\sigma_s$ at the subject level from data sampled at mid-cortical depth. Vertical black lines indicate the difference of zero. As expected, all participants show larger amplitudes (**A**) for GE- compared to SE-BOLD. The FWHM (**B**) similarly is broader for GE-BOLD for all but one participant (subject 5). Interestingly, this participant also had the smallest amplitude differences in **A**. Since the GE- and SE-BOLD data were acquired in separate sessions on different days, this discrepancy can have other reasons (e.g., differences in attentional states). Note that we used Bayesian hierarchical modeling to account for these residual differences at the population level. The baseline offset (**C**) shows no conclusive differences at the subject level. Comparisons of GE- and SE-BOLD parameter estimates at the population level are shown in ***Figure 8***.
